# CYP4F2-mediated ω-hydroxylation of 1-deoxysphingolipids reveals a new hepatic detoxification pathway

**DOI:** 10.64898/2026.05.11.724297

**Authors:** Adam Majcher, Essa M Saied, Zoltan Kutalik, Maftuna Shamshiddinova, Andreas J. Hülsmeier, Par Bjorklund, Elkhan Yusifov, Irina Alecu, Christoph Arenz, Thorsten Hornemann

**Affiliations:** Institute of Clinical Chemistry, University Hospital Zurich, Zurich, Switzerland; Humboldt Universität zu Berlin, Berlin D-12489, Germany; University Center for Primary Care and Public Health, University of Lausanne, 1010 Lausanne, Switzerland; Swiss Institute of Bioinformatics, 1015 Lausanne, Switzerland; Department of Medicine Huddinge, Cardio Metabolic Unit, Karolinska Institutet, Stockholm, Sweden; Clinical Department of Endocrinology, Karolinska University Hospital Huddinge, Stockholm, Sweden

## Abstract

1-deoxysphingolipids (1-deoxySLs) are atypical, cytotoxic sphingolipids (SL) formed by the serine palmitoyltransferase through the alternative use of L-Alanine over its canonical substrate L-Serine. Elevated plasma levels of 1-deoxySLs have been implicated in metabolic and neurodegenerative diseases. Due to the missing C1 hydroxyl group, 1-deoxySLs cannot be converted into complex sphingolipids nor degraded via the canonical SL catabolic pathways. However, previous reports suggested a cytochrome P450 mediated ω-hydroxylation of 1-deoxySLs as a potential detoxification mechanism although the exacts downstream metabolism of these lipids remained unclear. We combined genome-wide association analysis with targeted lipid analysis to identify genes involved in 1-deoxySL metabolism. Functional validation was performed in cell culture models, enzyme assays, and through quantitative high-resolution mass spectrometry using isotope labelled synthetic standards.We identified a strong association between the CYP4F2 rs2108622 variant and plasma 1-deoxySL, implicating CYP4F2 is involved in 1-deoxySL metabolism. We demonstrated that CYP4F2 catalyzes the ω-hydroxylation of 1-deoxysphinganine, forming a previously uncharacterized hydroxylated sphingoid base. In liver cells, this metabolite was further metabolized via three distinct pathways: one forming the N-acyl, a second involving omega acylation and third resulting in omega carboxylation. All reactions generated a new spectrum of 1-deoxysphingolipids that are based on ω-hydroxylated 1-deoxySA as a precursor. The metabolic steps were confirmed by structural validation using synthetically prepared external standards. Importantly, ω-hydroxylation significantly attenuated the acute cytotoxicity of 1-deoxySLs in liver cells, indicating that this modification is the initiating step of a multi-branched metabolic clearance pathway. This study identifies CYP4F2 as a key enzyme initiating the hepatic clearance of atypical 1-deoxySLs, mitigating their cellular toxicity and revealing multiple downstream metabolic fates. Our findings highlight a previously unrecognized clearance mechanism for atypical sphingolipids with relevance to metabolic disease.

## Introduction

1-Deoxysphingolipids (1-deoxySLs) are unusual sphingolipids produced during *de novo* sphingolipid synthesis, when serine-palmitoyltransferase uses L-alanine instead of its normal substrate, L-serine. Due to the missing C1-hydroxyl group, 1-deoxySLs cannot be converted into complex sphingolipids nor phosphorylated, rendering them resistant for the canonical SL catabolism (1). An abnormally high production of these lipids has been associated with multiple neurological conditions, such as hereditary sensory and autonomic neuropathy type 1 (HSAN1), diabetic sensory neuropathy and chemotherapy-induced peripheral neuropathy, where they contribute directly to disease development and progression (2-11). Toxicity has been linked to mitochondrial dysfunction, ER stress, and axonal degeneration (12-15).

Alecu et al. first reported that 1-deoxySLs undergo ω-hydroxylation by cytochrome P450 enzymes, particularly CYP4F11, offering a potential route for metabolic conversion and detoxification (16). However, the enzymes involved in this pathway, the formed downstream metabolites, and the physiological significance of 1-deoxySL ω-hydroxylation remains poorly defined.

Cytochrome P450 (CYP) enzymes constitute a large family of heme-containing monooxygenases involved in the metabolism of drugs, fatty acids, eicosanoids, and various xenobiotics. In the liver, CYP enzymes are central to both detoxification and lipid homeostasis, catalyzing oxidative reactions such as ω- and ω-1 hydroxylation (17). Among these, the CYP4F subfamily, including CYP4F2, CYP4F3, and CYP4F11, mediates terminal hydroxylation of bioactive lipids such as leukotriene B4, prostaglandins, hydroxyeicosatetraenoic acids (HETEs), and very long-chain fatty acids (VLCFAs), marking them for inactivation and clearance (18).

CYP4F2 is one of the most abundantly expressed hepatic isoforms, with established roles in the ω-hydroxylation of arachidonic acid, tocopherols, and VLCFAs. Genetic polymorphisms in CYP4F2, such as rs2108622, are known to affect enzyme activity and have been associated with inter-individual variability in warfarin response, vitamin E metabolism, and cardiovascular disease risk (19). While its function in fatty acid oxidation is well characterized, the role of CYP4F2 in sphingolipid metabolism has remained unexplored.

Genome-wide association studies (GWAS), particularly when combined with metabolite profiling, offer a powerful approach to identify genes involved in lipid metabolism. Integrative lipidomics GWAS have identified key regulators of plasma lipid traits, including phospholipids, ceramides, and acylcarnitines (20). In sphingolipid metabolism, variants in genes such as SPTLC3, SGPP1, DEGS1 and FADS3 have been linked to the abundance of specific lipid species (21, 22). In this study, we applied this approach to the CoLaus cohort, comprising of about 1,000 genotyped individuals with corresponding plasma lipid profiles acquired through targeted LC-MS/MS. By focusing on the plasma ratio of 1-deoxySA to 1-deoxySO, a proxy for 1-deoxySL turnover, we identified a genome-wide significant association with the CYP4F2 rs2108622 variant, implicating CYP4F2 in the metabolism of these atypical sphingolipids.

Building on these findings, we demonstrate that CYP4F2 catalyzes the ω-hydroxylation of 1-deoxySA, producing ω-hydroxy-1-deoxySA, a previously undescribed metabolite confirmed using synthetic standards. Functional studies in hepatic cells reveal that ω-OH-1-deoxySA is further metabolized into multiple novel lipid species, including ceramide-like structures, ω-O-acylated long-chain bases, and tritailed ceramides. These transformations involve both ceramide synthase-dependent and -independent steps, suggesting the existence of a broader detoxification clearance network for 1-deoxySLs.

Altogether, our findings identify CYP4F2 as a key enzyme mediating ω-hydroxylation and metabolic clearance of cytotoxic 1-deoxySLs and uncover a series of novel metabolic intermediates with potential physiological relevance.

## Results

### GWAS study reveals association of CYP4F2 with 1-deoxySL metabolism

Genome-wide association studies (GWAS), particularly when combined with metabolite profiling, offer a powerful approach(20) to identify genes involved in metabolism *(21, 22)* To identify genes involved in the conversion of 1-deoxySL, we analyzed single nuclear polymorphism (SNP) data from 1,100 individuals of the CoLaus cohort (7). Targeted long chain base (LCB) profiling was performed on acid-hydrolyzed plasma extracts to quantify the total levels of 1-deoxysphinganine (m18:0) and 1-deoxysphingosine (m18:1) from the same patients (Figure 1A).

**Figure 1:**
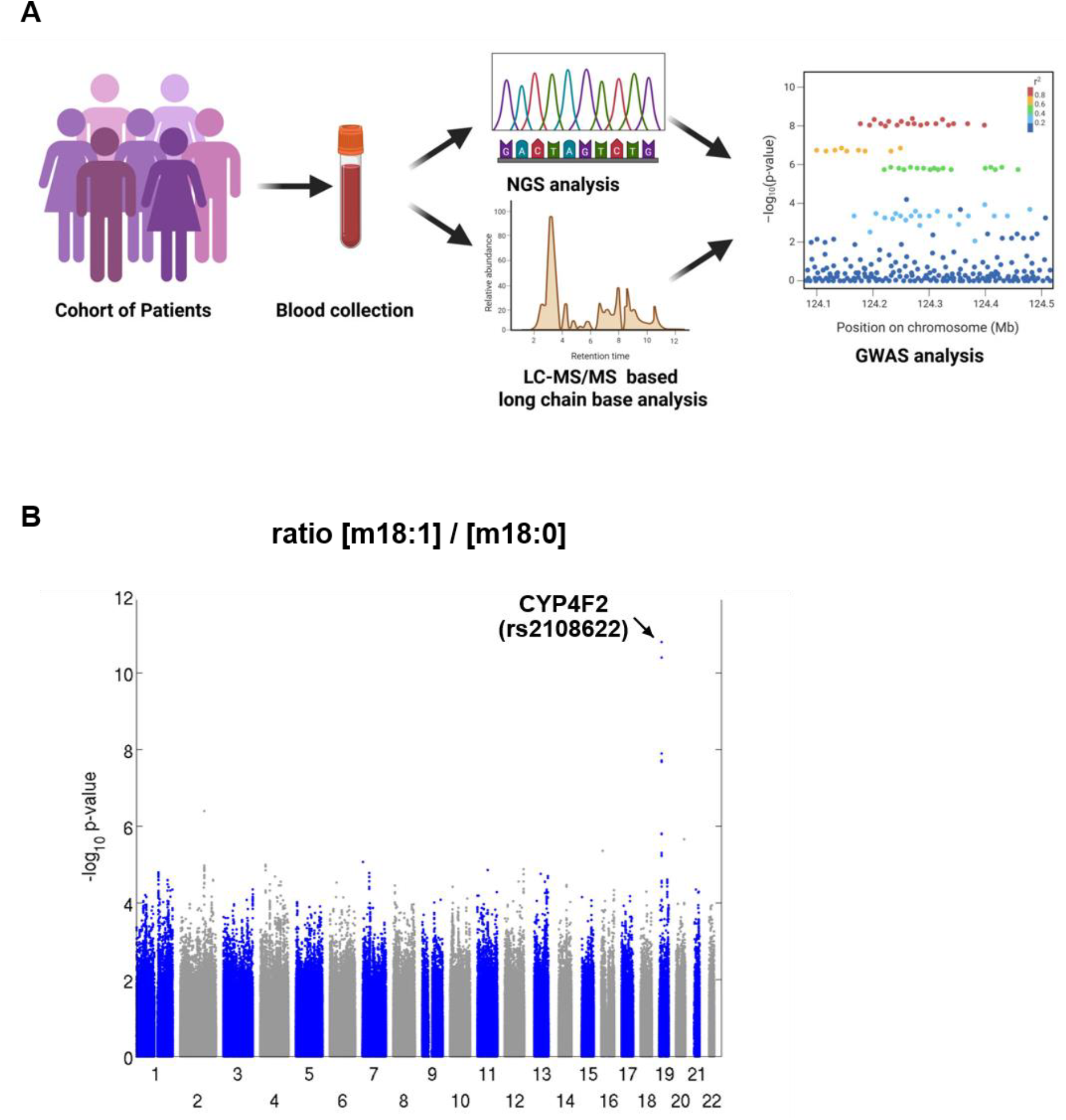
**(A)** Overview of the study design. Blood and plasma samples from 1,100 individuals of the CoLaus cohort were used for genome-wide SNP genotyping and targeted long chain base (LCB) profiling. Total LCB levels were measured in plasma samples after acid hydrolysis by LC-MS/MS. **(B)** Manhattan plot showing genome-wide association results for the ratio of the 1-deoxySO/1-deoxySA in plasma. The SNP rs2108622 in CYP4F2 showed a genome-wide significant association (p = 1.57 × 10^−11^), suggesting a role for CYP4F2 in 1-deoxySL metabolism.

The analysis revealed a significant genome-wide association (p = 1.57 × 10^−11^) between the SNP rs2108622 (Chr.19, Pos.15851431) and the [m18:1]/[m18:1] (Figure 1B). SNP rs2108622 is located in the CYP4F2 locus. Members of the CYP4F family have been previously suggested to be involved in the oxidative metabolism of 1-deoxySLs (16).

Previous studies have identified CYP4F2 as a hydroxylase of very long-chain fatty acids and bioactive lipids (18). Given the results from the GWAS and its role in lipid metabolism, we hypothesized that CYP4F2 might catalyze the hydroxylation of 1-deoxySL (Figure 2C). To test this, we treated HEK293 cells overexpressing CYP4F2 with isotope-labeled d3-1-deoxysphinganine (d3-1-deoxySA), followed by acid hydrolysis to release the free LCB, and subsequently performed high-resolution LC-MS/MS analysis. Extracted ion chromatograms (XICs) for the expected mass of m/z 305 [M+H^+^] revealed two isobaric peaks (Figure 2A): one at RT = 7.9 min and a second at RT = 8.3 min. The earlier-eluting metabolite corresponded to the hydroxylated product previously reported in MEF cells (16), whereas the later-eluting species had not been observed in MEF cells.

**Figure 2:**
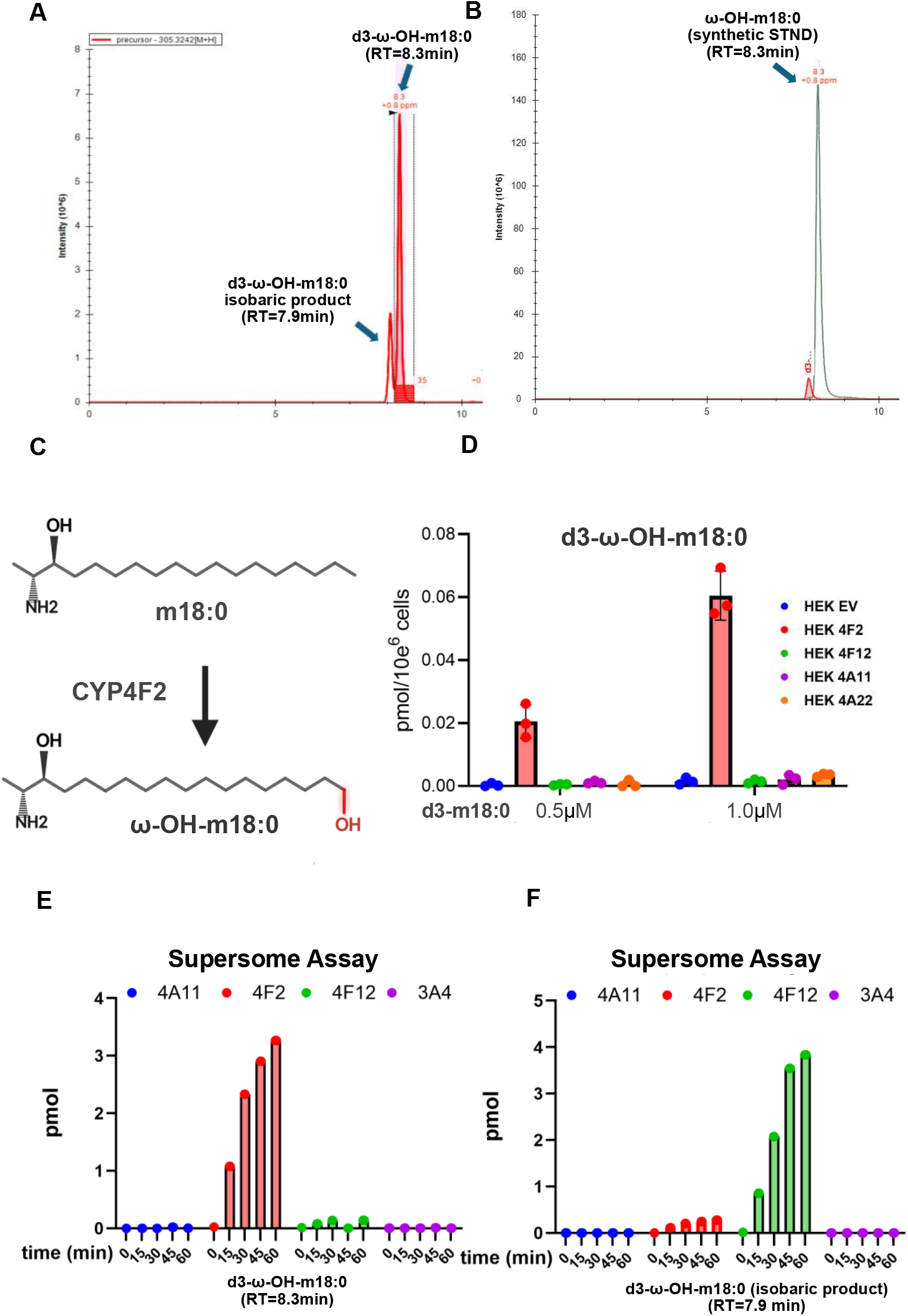
**(A)** Extracted ion chromatogram (XIC) of m/z 305 [M+H^+^], corresponding to d3-labeled hydroxylated 1-deoxySA, from HEK293 cells overexpressing CYP4F2 and treated with d3-1-deoxySA (1 µM, 16 h) after acid hydrolysis. Two isobaric peaks were detected: an earlier-eluting unknown hydroxylated isomer (RT = 7.9 min), and a later-eluting product (RT = 8.3 min) identified as d3-ω-hydroxy-1-deoxySA. **(B)** XICs of d3- and unlabeled hydroxylated 1-deoxySA (m/z 305 and 302 [M+H^+^]) from MEF cells treated with d3-1-deoxySA (1 µM, 16 h). Spiking a synthetic ω-OH-1-deoxySA confirmed that the peak at RT = 8.3 min corresponds to ω-hydroxylated 1-deoxySA, **C** Schematic illustrating the proposed CYP4F2-mediated ω-hydroxylation of 1-deoxySA. **D** Quantification of d3-ω-OH-1-deoxySA in HEK293 cells overexpressing different CYP isoforms (CYP4F2, red, CYP4F12, green, CYP4A11, purple, CYP4A22, orange, or empty vector, blue) and treated with d3-1-deoxySA (0.5 or 1 µM, 16 h).

To determine the identity of the second peak, we used MEF cell extracts spiked with synthetic ω-hydroxy-1-deoxySA (23). The spiked standard matched the retention time of the second peak at RT = 8.3 min (Figure 2B), confirming that this peak corresponds to ω-OH-1-deoxySA,a novel metabolite not previously described. The first peak at RT = 7.9 min has been reported as a sphingoid base but remains structurally uncharacterized.

To assess whether other CYP4F isoforms also contribute to 1-deoxySL metabolism, we compared the activity of various CYP isoforms in a commercial supersome *in vitro* assay (Figure 2C). Extracts from Sf9 insect cells heterologously expressing human CYP enzymes were spiked with d3-1-deoxySA, and the resulting metabolites were analyzed by LC-MS. Consistent with our cellular results, CYP4F2 specifically converted 1-deoxySA into ω-OH-1-deoxySA over time (Figure 2E). Interestingly, in CYP4F12-containing extracts, 1-deoxySA was instead converted into the isomer with RT = 7.9 min (Figure 2F).

These findings were further confirmed in HEK293 cells overexpressing CYP4F2, CYP4F12, CYP4A11, or CYP4A22 (Figure 2D). Formation of ω-OH-1-deoxySA was observed exclusively in CYP4F2-expressing cells, while CYP4F12, CYP4A11, and CYP4A22 showed no such activity (Figure 2D).

Together, these data demonstrate that CYP4F2 is the specific ω-hydroxylase responsible for the formation of ω-OH-1-deoxySA, whereas the previously described isomeric hydroxylated metabolite (RT = 7.9 min) appears to be associated with CYP4F12 but remains structurally uncharacterized

Only CYP4F2 induced ω-hydroxylation of 1-deoxySA. Time-course of d3-ω-OH-1-deoxySA formation in suprasome-based enzymatic assays with isolated CYP enzymes (CYP4F2, red, CYP4F12, green, CYP4A11, blue, CYP3A4, purple), confirming selective ω-hydroxylation by CYP4F2. **F** Time-course of the unknown d3-(OH)-1-deoxySA isomer (RT = 7.9 min) in suprasome assays, showing it is formed exclusively by CYP4F12. All long-chain bases were quantified by high-resolution LC-MS/MS and normalized to internal standards and total cell number.

### ω-Hydroxylation of 1-deoxySL as a detoxification mechanism

Cytochrome P450 enzymes, including the CYP4F family, are dominantly expressed in the liver, raising the question of whether the ω-hydroxylation of 1-deoxySA serves as a metabolic clearance mechanism. We therefore used Huh7 hepatoma cells to compare the cytotoxicity of the LCBs. Cells were treated with either 1-deoxySA, ω-OH-1-deoxySA, or 1-deoxySA in combination with the pan CYP4F inhibitor HET0016. Using the CellTiter-Glo viability assay, we observed that ω-OH-1-deoxySA was significantly less cytotoxic than 1-deoxySA (Figure 3A). Furthermore, inhibition of CYP4F activity increased 1-deoxySA toxicity, supporting the role of ω-hydroxylation in cellular protection.

**Figure 3:**
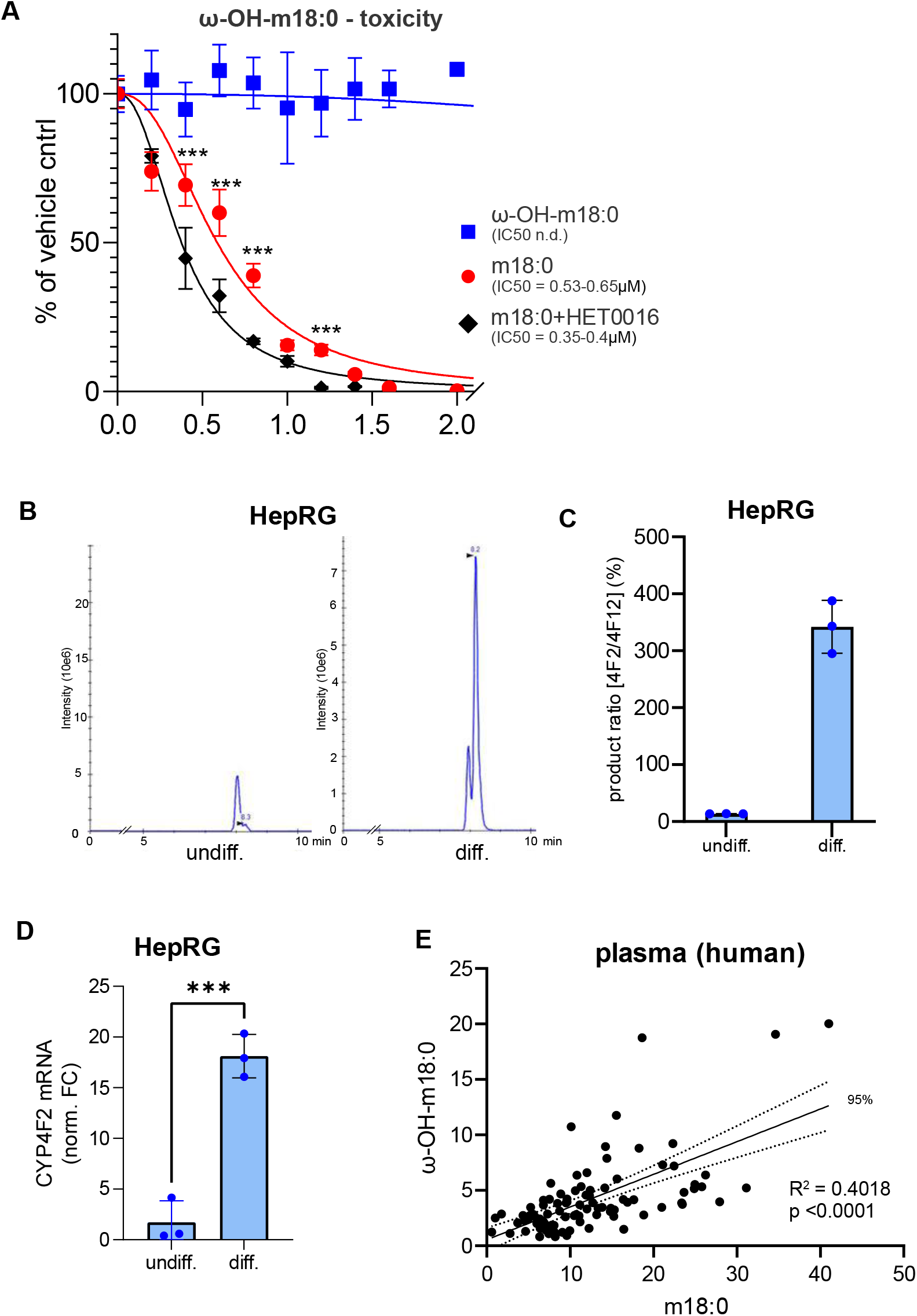
**(A)** Cell viability of Huh7 liver cells treated with 1-deoxySA (1 µM, red), synthetic ω-OH-1-deoxySA (1 µM, blue), or 1-deoxySA co-treated with the CYP4F-specific inhibitor HET0016 (2 µM, black). Cytotoxicity was assessed using the CellTiter-Glo assay and normalized to vehicle-treated controls. ω-OH-1-deoxySA showed significantly educed toxicity, while CYP4F inhibition enhanced 1-deoxySA-induced cytotoxicity. **(B)** qPCR analysis of CYP4F2 expression in undifferentiated vs. fully differentiated HepaRG cells.**(C, D)** Extracted ion chromatograms (XICs) of m/z 305 [M+H^+^] corresponding to d3-ω-OH-1-deoxySA in total long-chain base extracts from undifferentiated **(C)** and differentiated **(D)** HepaRG cells treated with d3-1-deoxySA (1 µM, 16 h). Differentiated cells showed markedly increased ω-hydroxylation, indicating an inducible hepatic clearance mechanism. **(E)** Correlation between plasma ω-OH-1-deoxySA and 1-deoxySA in human plasma (N = 93).

To investigate whether ω-hydroxylation is associated with hepatocyte maturity, we used the hepatocellular carcinoma line HepaRG that can develop into both hepatocyte and biliary epithelial-like cells. When differentiated into hepatocytes, they display many characteristics of primary human hepatocytes, including expression and activity of CYP450 enzymes, drug transporters and the synthesis of plasma proteins (e.g., albumin). We therefore treated both undifferentiated and differentiated HepaRG cells with d3-1-deoxySA and analyzed the formed downstream metabolites by LC-MS/MS. Conversion of 1-deoxySA into d3-ω-OH-1-deoxySA was largely absent in undifferentiated HepaRG cells but readily detectable in differentiated cells (Figure 3B,C), indicating that ω-hydroxylation is a feature of functionally mature HepaRG hepatocytes. Consistent with this maturation, qPCR analysis confirmed a strong upregulation of CYP4F2 expression in differentiated compared to undifferentiated cells (Figure 3D).

Finally, to determine whether this pathway is active in humans, we analyzed plasma samples from 93 individuals using targeted LC-MS/MS. We observed a significant positive correlation between 1-deoxySA and ω-OH-1-deoxySA levels (Figure 3E), suggesting a shared metabolic origin and further supporting that ω-hydroxylation is a physiologically relevant, liver-associated route of 1-deoxySL clearance.

### Discovery and identification of Omega hydroxy 1-deoxyDHCeramides

To determine whether ω-hydroxylation of 1-deoxySA precedes N-acylation (Figure 4A), we treated CYP4F2 overexpressing HEK293 cells with d3-1-deoxySA in the presence or absence of the pan-ceramide synthase (CerS) inhibitor Fumonisin B1 (FB1). The addition of FB1 led to a marked accumulation of d3-ω-OH-1-deoxySA (Figure 4B), indicating that ω-hydroxylation occurs primarily on the free long-chain base and precedes N-acylation by CerS.

**Figure 4:**
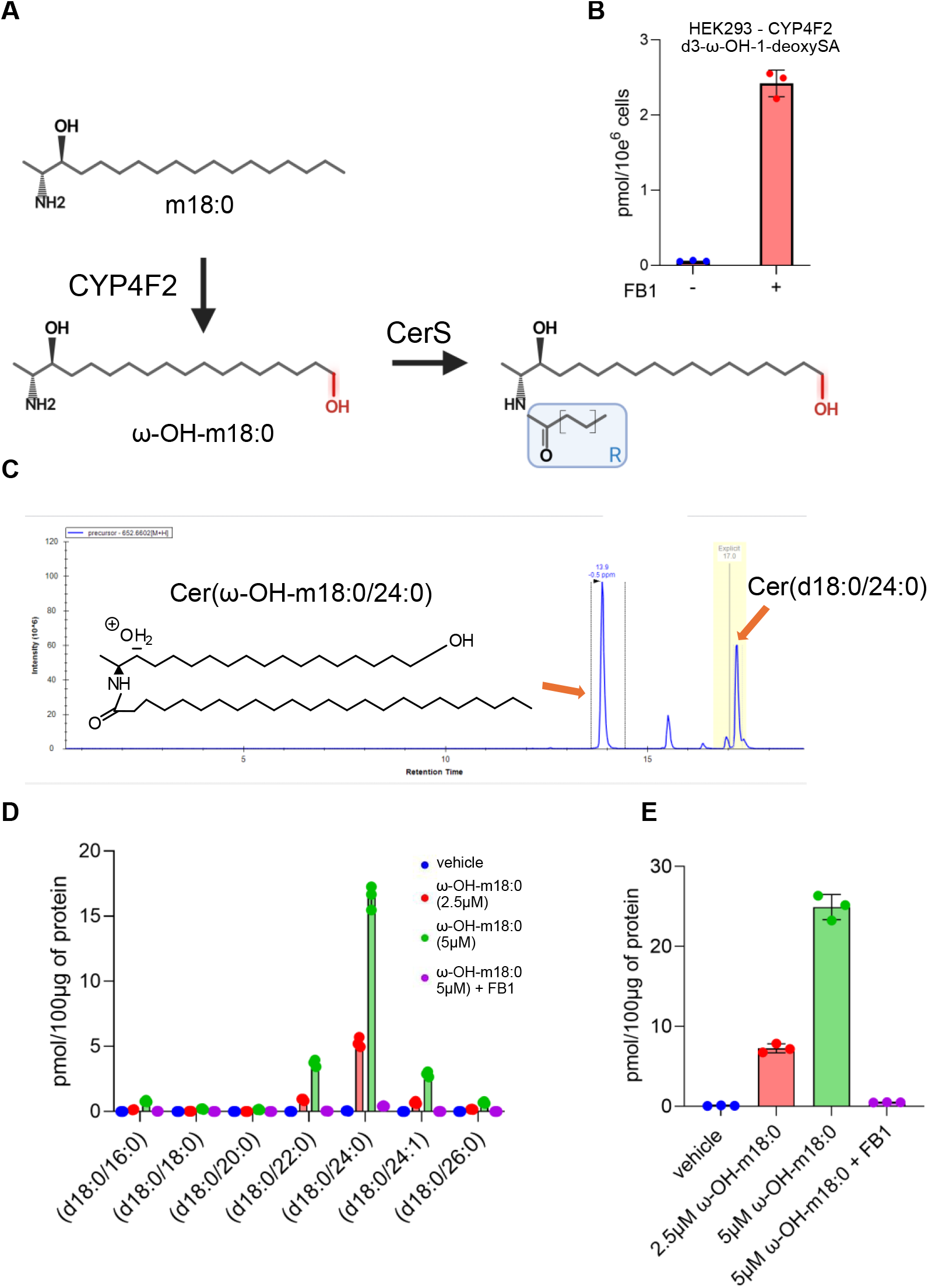
**(A)** Schematic overview of the proposed metabolic pathway downstream of CYP4F2-mediated ω-hydroxylation of 1-deoxySA. We hypothesize that ω-OH-1-deoxySA (ω-OH-m18:0) serves as a substrate for ceramide synthases (CerS), forming a novel class of ω-hydroxylated 1-deoxy-dihydroceramides. **(B)** Quantification of d3-ω-OH-1-deoxySA in HEK293 cells overexpressing CYP4F2 and treated with d3-1-deoxySA (1 µM, 16 h), with or without the CerS inhibitor fumonisin B1 (FB1, 35 µM). Accumulation of the free base under CerS inhibition indicates that hydroxylation precedes N-acylation and supports its role as a CerS substrate. **(C)** Extracted ion chromatogram (XIC) of m/z 652 [M+H^+^], corresponding to Cer(ω-OH-m18:0/24:0), in lipid extracts from Huh7 cells treated with synthetic ω-OH-1-deoxySA (5 µM, 16 h). The earlier retention time relative to canonical Cer(d18:0/24:0) reflects altered polarity from the ω-hydroxyl group. Structural identity was confirmed by MS^2^ fragmentation, showing diagnostic fragments of both the ω-hydroxylated sphingoid base and N-acyl amide. **(D)** Quantification of individual N-acyl-ω-OH-m18:0 species and () Cumulative abundance of all N-acyl-ω-OH-1-deoxySA in Huh7 cells treated with ω-OH-1-deoxySA (2.5 µM or 5 µM, 16 h), with or without FB1 (35 µM). Ceramide formation was fully blocked by CerS inhibition, confirming CerS-dependent synthesis. Lipids were quantified using high-resolution untargeted LC-MS/MS lipidomics and normalized to internal standard and total protein content.

Furthermore, we hypothesized that also ω-OH-1-deoxySA may serve as a substrate for CerS, thereby formation of previously non described classes of ω-hydroxylated 1-deoxy-(dihydro)ceramides.

To understand the full network of 1-deoxySL downstream metabolism, we treated Huh7 cells with ω-OH-1-deoxySA and characterized the metabolic downstream product with m/z 652 [M+H^+^], corresponding to a N-acyl-ω-OH-1-deoxySA that equals the most abundant ω-OH-deoxyDHCer (m18:0-OH/24:0). ω-OH-Ceramide species eluted significantly earlier than the canonical Cer(d18:0/24:0), due to the reduced hydrophobicity conferred by the ω-hydroxyl group. Structural confirmation of N-acyl-ω-OH-1-deoxySA was achieved via MS^2^ fragmentation, which resulted in the characteristic fragments of the ω-hydroxylated sphingoid base and the corresponding amide moiety, both consistent with the proposed structure.

In addition, supplementation of Huh7 cells with ω-OH-1-deoxySA led to a dose-dependent accumulation of multiple N-acyl-ω-OH-1-deoxySA species (Figure 4D), characterized by a CerS2-associated C22–C24 N-acyl chain profile. Co-treatment with FB1 completely abolished the formation of these metabolites (Figure 4E), confirming that ω-OH-deoxyDHCer serves as a substrate for CerS.

### Identification of omega-O-Acylated derivatives of ω-OH-1-deoxySphinganine

Given the presence of a terminal ω-hydroxyl group on ω-OH-1-deoxySA, we hypothesized that this moiety may also serve as a substrate for enzymatic esterification by an as-yet-unidentified ω-O-acyltransferase. To investigate this possibility, we analyzed lipid extracts from Huh7 cells treated with synthetic ω-OH-1-deoxySA using high-resolution untargeted LC-MS/MS. Besides the above mentioned ω-OH-1-deoxySA species, we also identified a distinct class carrying an O-acyl group at the ω-position, which we designated as ω-O-acyl-m18:0 long-chain bases (LCBs).

The structural identity of ω-O-acyl-m18:0 LCBs was confirmed by a unique MS^2^ fragmentation profiles. Unlike canonical sphingoid base fragments that retain the amine group and exhibit an even nominal mass, these spectra were characterized by ester-linked fatty acyl product ions lacking nitrogen (resulting in an odd nominal mass, e.g., m/z 239). The mass of these diagnostic acyl fragments shifted precisely in accordance with the length of the esterified acyl chain. Representative spectra of d18:0/0:0/O-16:0 and d18:0/0:0/O-14:0 species revealed a consistent sphingoid base backbone and diagnostic ester fragment patterns, including a characteristic +28 Da (C_2_H_4_) shift (Figure 5A–B).

**Figure 5:**
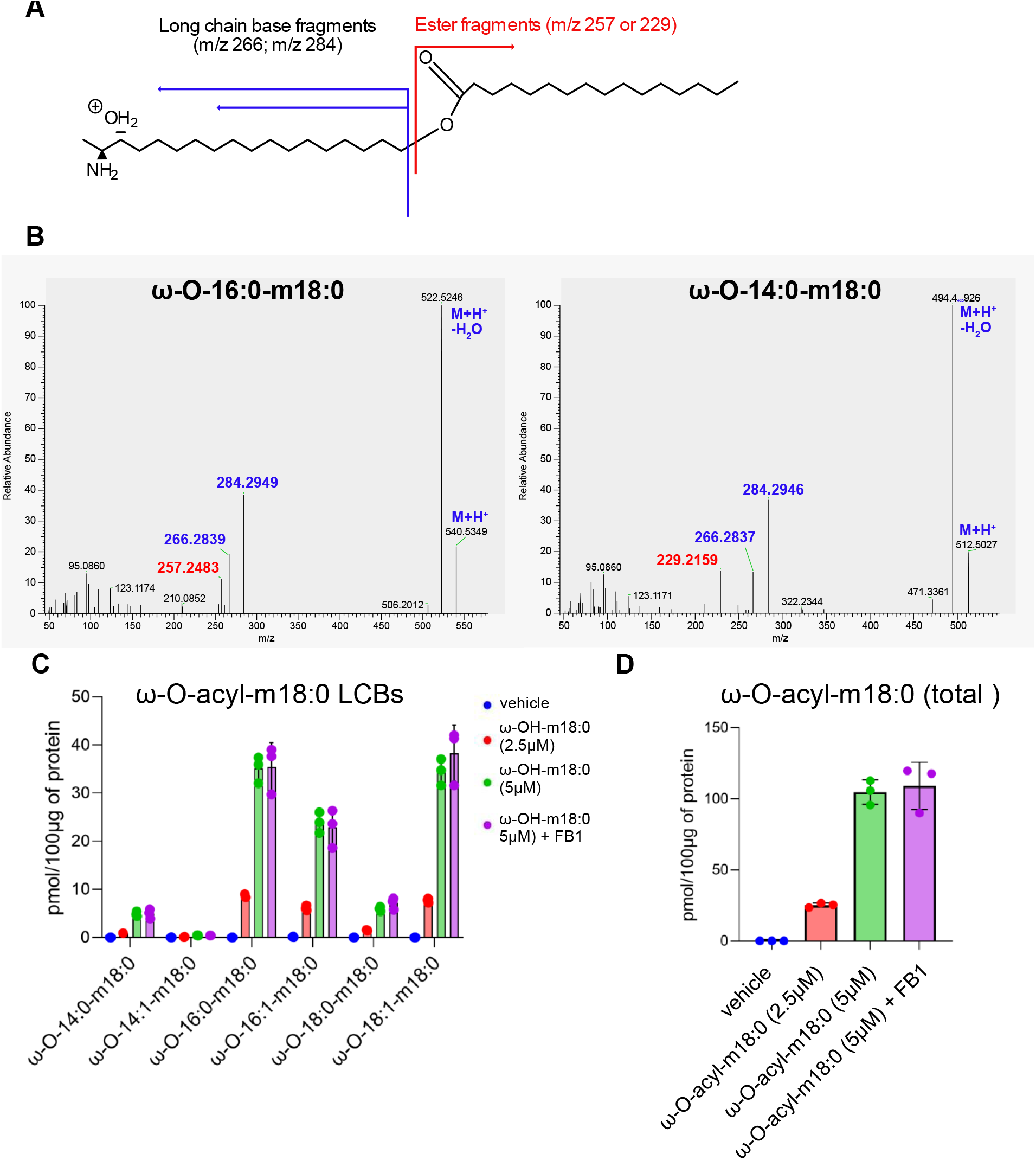
**(A)** Schematic representation of the chemical structure and characteristic MS^2^ fragmentation pattern of ω-O-acyl-m18:0 long-chain bases (LCBs), a novel class of ω-O-acylated sphingoid bases. These lipids were identified by diagnostic ester-linked fatty acyl product ions. In contrast to canonical nitrogen-containing sphingoid base fragments (even m/z values), these nitrogen-free ester fragments presented with an odd m/z value and exhibited mass shifts corresponding precisely to the length of the esterified acyl chain. For detailed nomenclature, see Table 1. **(B)** Representative MS^2^ spectra of two ω-O-acyl-m18:0 LCB species (ω-O-16:0-m18:0 and ω-O-14:0-m18:0). The ester fragments show a +28 Da (C_2_H_4_) shift between species, while the long-chain base fragment remains constant, confirming ω-O-acylation. **(C)** Quantification of individual ω-O-acyl-m18:0 LCB species and (**D)** Total ω-O-acyl-m18:0 LCB levels in Huh7 cells treated with ω-OH-m18:0 (2.5 or 5 µM, 16 h), with or without the ceramide synthase inhibitor fumonisin B1 (FB1; 5 µM and 35 µM). Formation of ω-O-acyl-m18:0 LCBs was FB1-insensitive, suggesting a CerS-independent pathway. Lipids were analyzed using high-resolution untargeted LC-MS/MS and normalized to internal standard and total protein content.

Quantification of ω-O-acyl-m18:0 LCBs showed a prominent use of medium to long chain fatty acids, with C14:0, C16:0, C16:1, C18:0, and C18:1 being the most abundant species (Figure 5C). We did not observe the formation of ω-O-acyl-m18:0 metabolites with ω-O-acyl chains ≥ C20.

**Table 1:** Nomenclature of the 1-deoxySphingolipid metabolites.

| Structural Description | Proposed Shorthand | Example Species | LIPID MAPS Systematic Notation |
| --- | --- | --- | --- |
| ω-OH-1-deoxy sphinganine | ω-OH-m18:0 | ω-OH-m18:0 | SPB 18:0;3OH;18OH |
| ω-OH-1-deoxy sphingosine | ω-OH-m18:1 | ω-OH-m18:1 | SPB 18:1;3OH;18OH |
| N-acyl-ω-OH-1-deoxy sphinganine | Cer(ω-OH-m18:0/xx:y) | Cer(ω-OH-m18:0/24:0) | Cer 18:0;3OH;18OH/24:0 |
| N-acyl-ω-OH-1-deoxy sphingosine | Cer(ω-OH-m18:1/xx:y) | Cer(ω-OH-m18:1/24:0) | Cer 18:1;3OH;18OH/24:0 |
| ω-O-acyl-1-deoxy sphinganine | ω-O-acyl-m18:0 | ω-O-16:0-m18:0 | SPB 18:0;3OH;18O(16:0) |
| ω-O-acyl-1-deoxy sphingosine | ω-O-acyl-m18:1 | ω-O-16:0-m18:1 | SPB 18:1;3OH;18O(16:0) |
| N-acyl-ω-O-acyl-1-deoxy sphinganine | Cer(ω-O-acyl-m18:0/xx:y) | Cer(ω-O-16:0-m18:0/24:0) | Cer 18:0;3OH;18O(16:0)/24:0 |
| N-acyl-ω-O-acyl-1-deoxy sphingosine | Cer(ω-O-acyl-m18:1/xx:y) | Cer(ω-O-16:0-m18:1/24:0) | Cer 18:1;3OH;18O(16:0)/24:0 |
| ω-carboxy-1-deoxy sphinganine | ω-COOH-m18:0 | ω-COOH-m18:0 | SPB 18:0;3OH;18COOH |
| ω-carboxy-1-deoxy sphingosine | ω-COOH-m18:1 | ω-COOH-m18:1 | SPB 18:1;3OH;18COOH |
| N-acyl-ω-carboxy-1-deoxy sphinganine | Cer(ω-COOH-m18:0/xx:y) | Cer(ω-COOH-m18:0/24:0) | Cer 18:0;3OH;18COOH/24:0 |
| N-acyl-ω-carboxy-1-deoxy sphingosine | Cer(ω-COOH-m18:1/xx:y) | Cer(ω-COOH-m18:1/24:0) | Cer 18:1;3OH;18COOH/24:0 |

The formation of ω-O-acyl-m18:0 LCBs was not altered in presence of the CerS inhibitor FB1 (Figure 5D), indicating that their biosynthesis is independent of canonical ceramide synthase activity. These findings suggest that ω-O-acylation is not mediated by CerS but another acyltransferase which identity remains to be elucidated.

The newly identified ω-hydroxylated and ω-O-acylated 1-deoxy-sphingolipids were named according to the adapted sphingolipid nomenclature outlined in Table 1, providing consistent annotation of these structurally distinct metabolite classes.

### Identification of tritailed 1-deoxy-Sphingolipid derivatives

Building on our identification of an ω-O-acyl-m18:0 long-chain bases, we hypothesized that the ω-O-acylated intermediates may also serve further as substrates for ceramide synthases, yielding a structurally distinct class of ω-O-acyl-1-deoxy-ceramides (Figure 6A). These lipids are defined as 1-deoxy-sphingoid base backbone, bearing both an N-acyl and an ω-O-acyl moiety. Their nomenclature and structural classification are provided in Table 1.

**Figure 6:**
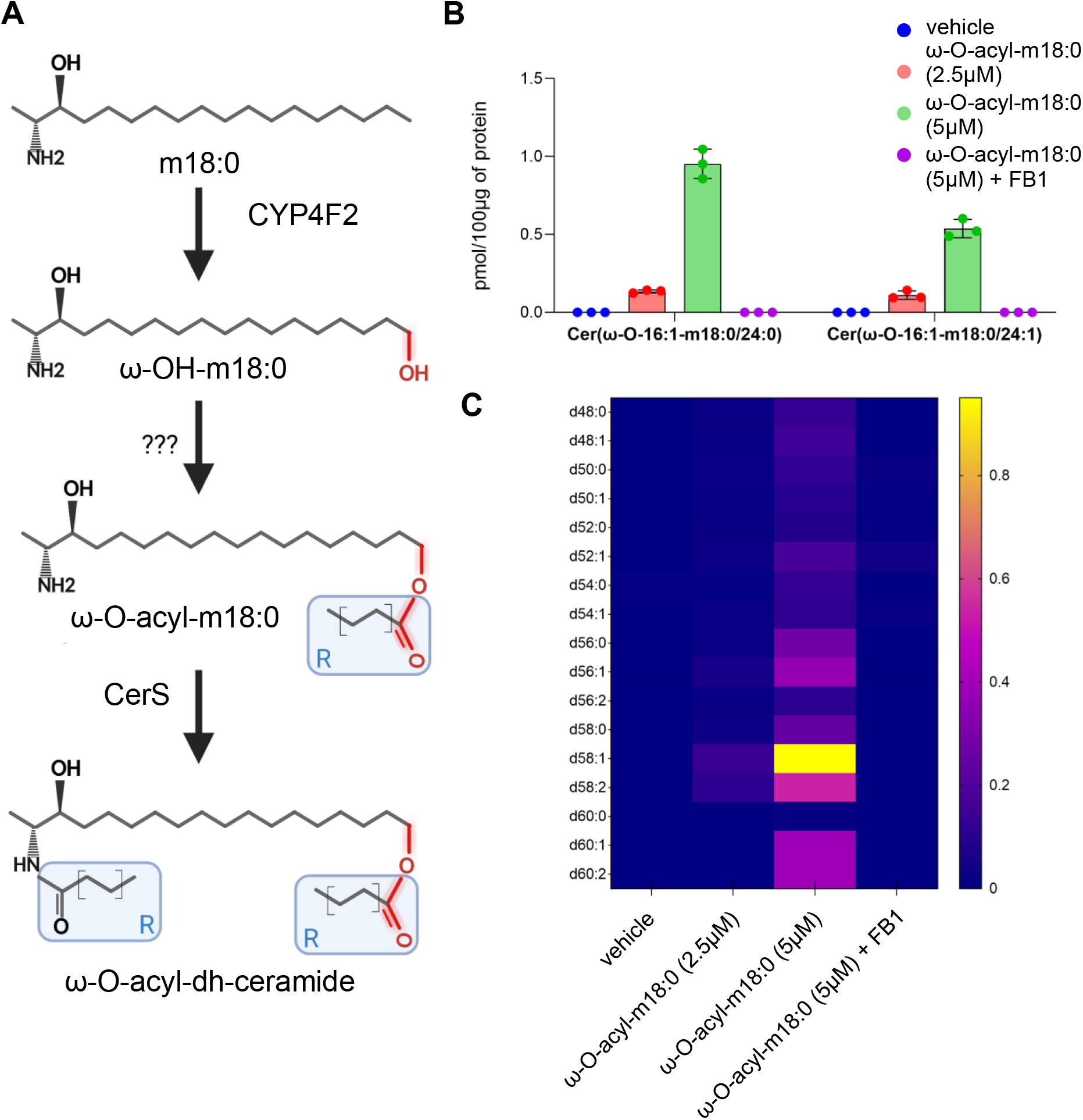
**(A)** Schematic illustrating the metabolic pathway following ω-hydroxylation of m18:0 by CYP4F2 and subsequent acylation of ω-OH-m18:0 by an unknown enzyme to form ω-O-acyl-m18:0. We hypothesize that ω-O-acyl-m18:0 serve as substrates for CerS, leading to the formation of a previously undescribed class of N-acyl-ω-O-acyl-1-deoxy-LCBs. For nomenclature details, refer to Table 1. **(B)** Quantification of two characterized ω-O-acyl-1-deoxy-ceramides at the subspecies level. **(C)** Quantification of additional potential ω-O-acyl-1-deoxy-ceramides in Huh7 cells treated with ω-OH-m18:0 (2.5 µM or 5 µM, 16 hours), with or without co-treatment with the CerS inhibitor FB1 (35 µM). These potential ω-O-acyl-1-deoxy-ceramide species were quantified at the lipid species level due to insufficient MS^2^ fragmentation data and lack of synthetic standards preventing full structural confirmation. Lipids were quantified using a high-resolution untargeted LC-MS/MS lipidomics workflow and normalized to internal standard and total protein content.

To evaluate this pathway, Huh7 cells were treated with synthetic ω-OH-1-deoxySA, and lipid extracts were analyzed by high-resolution untargeted LC-MS/MS. The presence of two ω-O-acyl-1-deoxy-dihydro-ceramide species were fully confirmed by MS^2^ fragmentation, demonstrating characteristic product ions consistent with both N- and O-acyl substituents (Figure 6B). Other putative ω-O-acyl-1-deoxy-dihydro-ceramide species were identified on parent ion level according to its calculated exact mass but could not full confirmed, due to limited fragmentation and the absence of synthetic standards (Figure 6C).

### Identification of ω-carboxy-1-deoxy-Sphinganine and ω-Carboxy-1-deoxy-dihydro-ceramides

We identified a subset of metabolites derived from ω-hydroxylated 1-deoxysphinganine (ω-OH-1-deoxySA) that displayed a +14 Da mass shift and distinct chromatographic retention times compared to ω-OH-1-deoxy-ceramides species. These features are consistent with an additional oxidation step, specifically ω-carboxylation, potentially representing a previously undescribed step of 1-deoxySL metabolism (Figure 7A).

**Figure 7:**
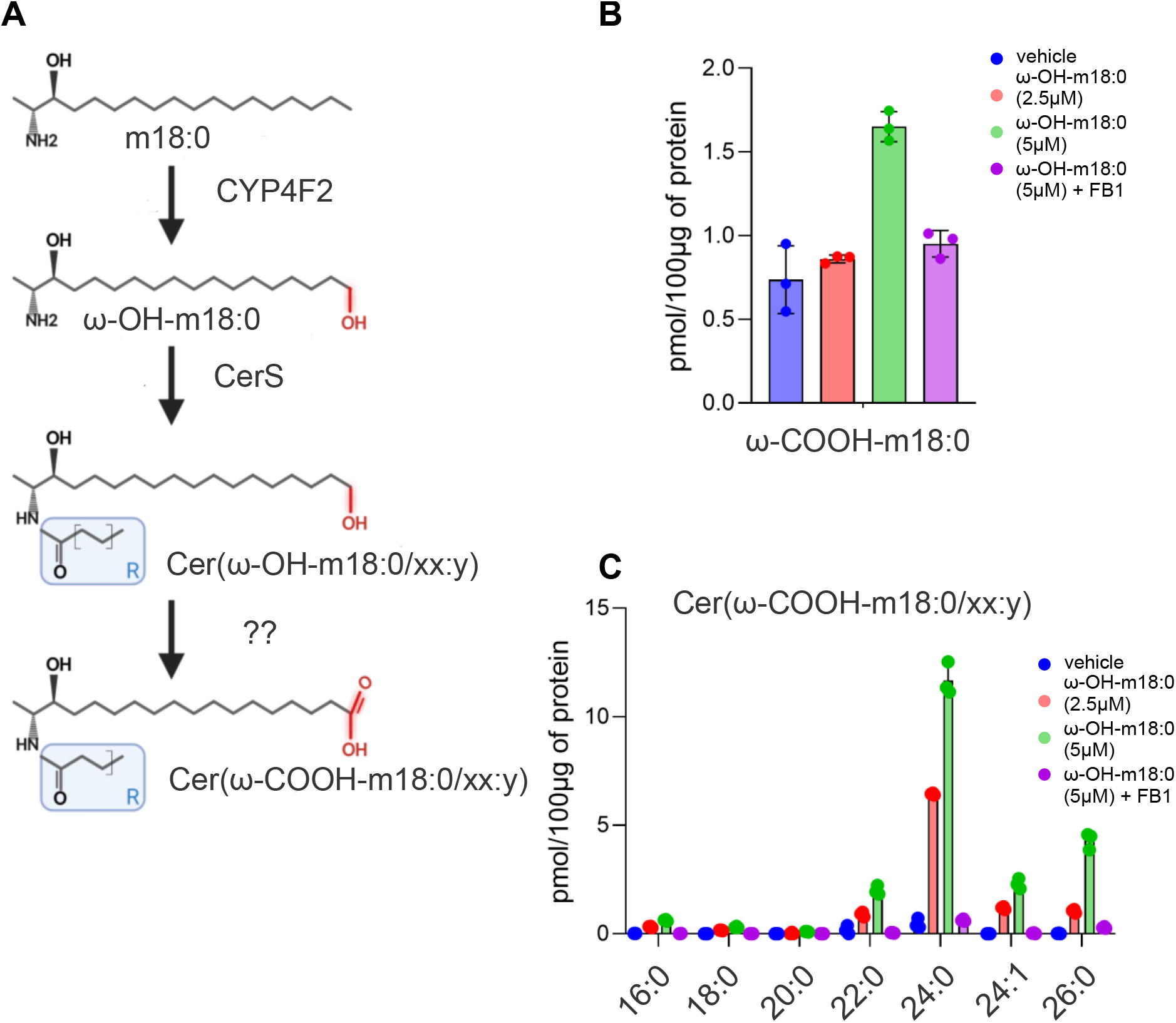
**(A)** Schematic illustrating the proposed hypothetical metabolic pathway following ω-hydroxylation of m18:0 by CYP4F2, N-acylation by CerS and subsequent oxidation of ω-OH-1-deoxy-dihydro-ceramide (Cer(ω-OH-m18:0/xx:y)) by an unknown enzyme to form ω-carboxylated 1-deoxy-dihydro-ceramide (Cer(ω-COOH-m18:0/xx:y)). For the detailed nomenclature please check Table 1. **(B)** Quantification of individual ω-COOH-1-deoxy-dihydro-ceramide species (vehicle, blue, 2.5 µM ω-OH-m18:0, red, 5 µM ω-OH-m18:0, green, 5 µM ω-OH-m18:0 + FB1, purple) and **(C)** free ω-COOH-m18:0 levels in Huh7 cells treated with ω-OH-m18:0 (2.5 or 5 µM, 16 h), with or without the ceramide synthase inhibitor fumonisin B1 (FB1; 35 µM, vehicle, blue, 2.5 µM ω-OH-m18:0, red, 5 µM ω-OH-m18:0, green, 5 µM ω-OH-m18:0 + FB1, purple). Lipids were analyzed using high-resolution untargeted LC-MS/MS and normalized to internal standard and total protein content.

Based on mass spectrometry and chromatographic behavior, we hypothesized that these metabolites represent ω-carboxylated derivatives, namely ω-COOH-1-deoxySA (ω-COOH-m18:0 LCB) and its N-acylated counterpart, ω-COOH-1-deoxy-dihydro-ceramide (Figure 7A). Formation of ω-COOH-1-deoxy-dihydro-ceramide was abolished by co-treatment with FB1 (Figure 7B, C), indicating that their biosynthesis depends on CerS activity.

Because long-chain bases exhibit non-specific MS^2^ fragmentation patterns, structural confirmation of ω-COOH-1-deoxySA by fragmentation alone was inconclusive. To proof the structural identity of the metabolite we chemically synthesized a ω-COOH-1-deoxySA reference standard (see Supplementary Methods and Supplementary Figure S1). Huh7 cells, were supplemented with either synthetic ω-OH-1-deoxySA or the ω-COOH-1-deoxySA and the sphingolipid profile analyzed by untargeted high-resolution LC-MS/MS (Figure 8A).

**Figure 8:**
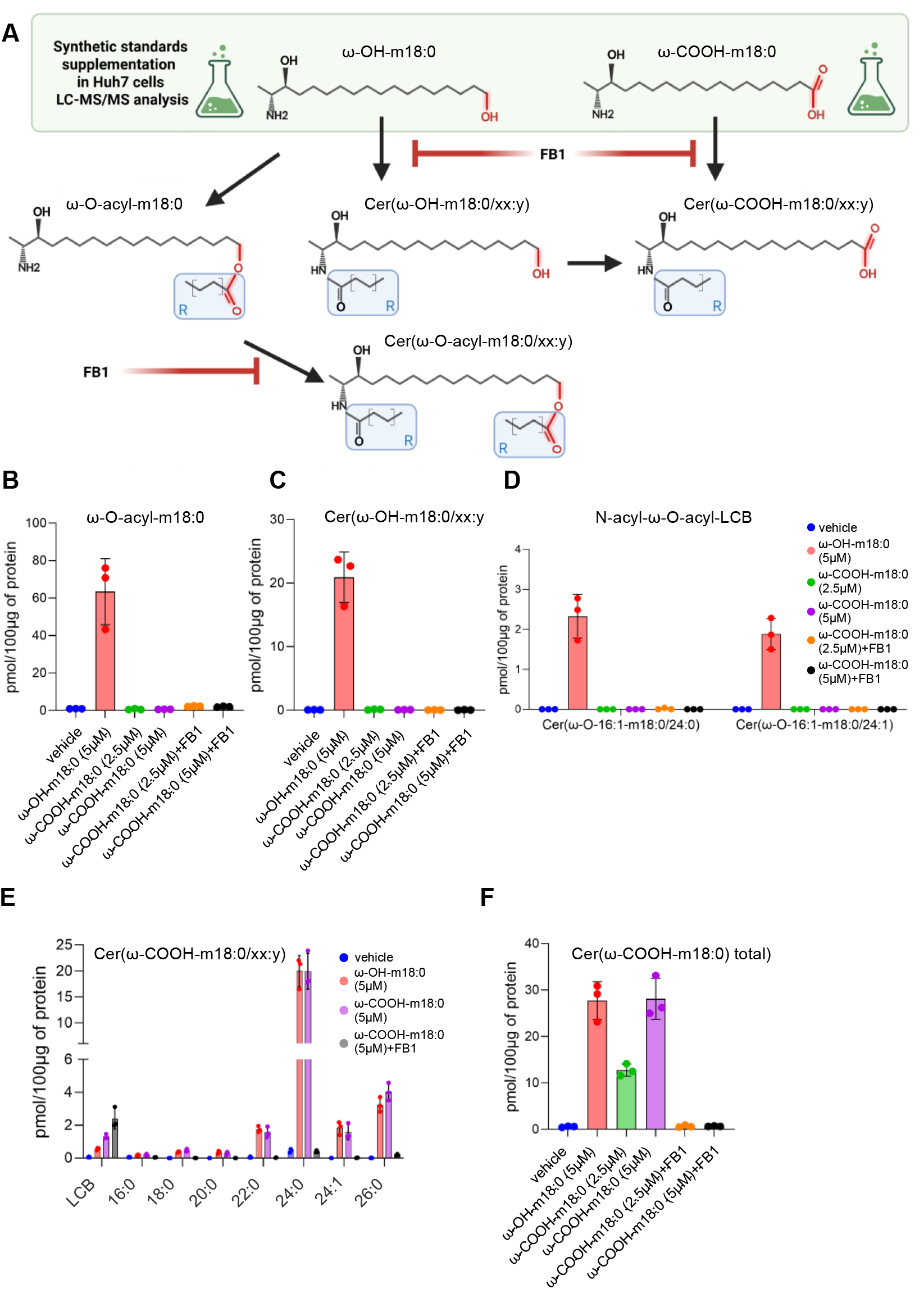
**(A)** Schematic illustrating the metabolic tracking of synthetic ω-OH-m18:0 and ω-COOH-m18:0 standards. Fumonisin B1 (FB1) was used as a ceramide synthase (CerS) inhibitor to prevent the N-acylation of the long-chain bases. **(B–F)** Huh7 cells were treated for 16 h with vehicle (blue), 5 µM ω-OH-m18:0 (red), 2.5 µM or 5 µM ω-COOH-m18:0 (purple) or co-treated with 5 µM ω-COOH-m18:0 and 35 µM FB1 (black). Bar graphs show the quantification of: **(B)** total ω-O-acyl-m18:0 LCBs; **(C)** total ω-OH-1-deoxy-dihydro-ceramides; **(D)** two identified tritailed N-acyl-ω-O-acyl-LCBs; () free ω-COOH-m18:0 LCB and individual ω-COOH-1-deoxy-dihydro-ceramide species; and **(F)** total ω-COOH-1-deoxy-dihydro-ceramides. Lipids were analyzed using high-resolution untargeted LC-MS/MS and normalized to an internal standard and total cellular protein content

Notably, only ω-OH-1-deoxySA, but not the ω-COOH-1-deoxySA standard, gave rise to the previously characterized downstream metabolites such as ω-O-acyl-m18:0 LCB, ω-COOH-1-deoxy-dihydro-ceramide and ω-O-acyl-dihydro-ceramide (Figure 8B–D). This demonstrates that these downstream species are metabolically derived from ω-OH-1-deoxySA rather than from ω-COOH-1-deoxySA, indicating that ω-carboxylation represents a distinct, divergent branch in 1-deoxySL metabolism.

Retention time alignment and MS feature comparison confirmed that the ω-carboxylated species are formed after ω-OH-1-deoxySA supplementation and are structurally identical to the synthetic ω-COOH-1-deoxySA as well as its downstream ceramide, ω-COOH-1-deoxy-dihydro-ceramide (Figure 8E, F). Moreover, FB1 treatment blocked the formation of ω-COOH-1-deoxy-dihydro-ceramide from the ω-carboxylic precursor, verifying that ω-COOH-1-deoxySA is also a substrate for CerS.

Together, these findings expand the known spectrum of 1-deoxySL metabolites to include species generated by a secondary ω-carboxylation step following CYP4F2-mediated ω-hydroxylation. This newly described carboxylated long-chain base adds further complexity and metabolic flexibility to the cellular metabolism of cytotoxic 1-deoxySLs.

## Discussion

1-Deoxysphingolipids (1-deoxySLs) are atypical, cytotoxic sphingolipid species synthesized by serine palmitoyltransferase (SPT) through the incorporation of L-alanine instead of L-serine. Their lack of a C1 hydroxyl group renders them resistant to canonical catabolic processes and prevents conversion into complex sphingolipids, leading to accumulation and toxicity (1). Pathologically elevated 1-deoxySLs have been linked to metabolic disorders, including T2D, HSAN1, and chemotherapy-induced neuropathies (2, 3, 5-11, 24, 25). Thus, identifying mechanisms of detoxification and metabolic clearance is essential.

Building on prior work by Alecu et al. (16), which first identified the CYP4F family -particularly CYP4F11 - as capable of hydroxylating 1-deoxySLs *in vitro*, our study establishes CYP4F2 as the dominant, physiologically relevant ω-hydroxylase in humans. While biochemical redundancy undoubtedly exists among CYP4F isoforms, our unbiased GWAS data reveals a genome-wide significant association between the CYP4F2 rs2108622 variant and altered plasma 1-deoxySL turnover. This strong human genetic signal provides powerful *in vivo* evidence that other isoforms, such as CYP4F11, cannot fully compensate for reduced CYP4F2 activity. Therefore, while our biochemical validation relies on overexpression models rather than selective knockout cells, the human population data firmly positions CYP4F2 as the primary, clinically relevant driver of this hepatic clearance pathway. Furthermore, our targeted high-resolution LC-MS/MS data expands upon previous findings by structurally confirming the specific formation of ω-OH-m18:0 (RT=8.3 min) using a synthetic standard, distinguishing it from other uncharacterized CYP-derived isomers.

Furthermore, our *in vitro* assays revealed a distinct, earlier-eluting (RT = 7.9 min) hydroxylated isomer produced exclusively by CYP4F12. While the exact regiochemistry of this minor isomer remains to be structurally elucidated, its discovery suggests that multiple CYP4F isoforms may engage 1-deoxySLs to produce a diverse array of positional isomers. Rather than a singular metabolic dead-end, this points to a highly complex, multi-isoform regulatory network. Characterizing this parallel CYP4F12-driven pathway, and the bioactivity of its resulting isomer, represents a compelling avenue for future research to fully map the metabolic fate of 1-deoxySLs.

CYP4F2 belongs to the CYP4F subfamily of cytochrome P450 monooxygenases, which are known for ω- and ω-1 hydroxylation of endogenous lipids, including arachidonic acid, leukotriene B4, prostaglandins, and HETEs. These modifications generally reduce bioactivity and facilitate clearance (18). CYP4F2 has also been implicated in warfarin resistance through its role in vitamin K ω-hydroxylation, which lowers hepatic vitamin K levels less efficiently in carriers of reduced-function variants, thereby contributing to inter-individual variability in warfarin dose requirements.

Our data extend the list of possible substrates showing that also deoxySL are metabolized by this enzyme. The mechanism mirrors previously established roles of CYP4F2 in hepatic toxin clearance.

In this study, using a large human cohort with matched lipidomic and genomic data, to identify a genome-wide significant association between the CYP4F2 rs2108622 variant and the 1-deoxySA/1-deoxySO ratio. A similar approach has been used to identify FADS3 being involved in 1-deoxySL metabolism. In contrast to CYP4F2, FADS3 introduces a 14Z double bond to convert 1-deoxySA into 1-deoxySO (m18:1), a less toxic 1-deoxySL derivative (21).

While ω-hydroxylation by CYP4F2 leads to metabolic clearance via formation of less toxic hydroxylated species, we further show that this initial modification is followed by subsequent metabolic conversion. In Huh7 cells, ω-OH-1-deoxySA was incorporated by CerS into novel ceramide-like and esterified 1-deoxySL species. Similarly, also other LCBs such as 3-ketodihydrosphinganine is N-acylated and mutations in KDSR result in progressive symmetric erythrokeratoderma (PSEK) a rare inherited keratinization disorder of the skin cause by an increased formation of 3-keto-Ceraminde (26).

We also showed that ω-OH-1-deoxySA is further O-acylated forming a new class of O-acylated 1-deoxy-sphingoid bases. These structures differ from canonical ceramides due to their tritailed structure. Notably, the O-acylation process was insensitive to fumonisin B1, suggesting that ceramide synthases (CerS) are not involved and this reaction is mediated by a yet unidentified enzyme.

In addition to O-acylation, we identified ω-carboxylation as a further downstream modification of ω-OH-deoxySL. This process appears to convert ω-OH-1-deoxySA into ω-COOH-1-deoxySA and also in its N-acylated derivative ω-COOH-1-deoxy-dihydro-ceramide. The identity of ω-COOH-1-deoxySA was confirmed by comparison to a chemically synthesized ω-COOH-1-deoxySA standard. The formation of ω-COOH-1-deoxy-dihydro-ceramide was abrogated by by FB1 treatment, indicating a CerS-dependent carboxylation reaction. While structural confirmation by MS^2^ fragmentation alone was limited due to a non-specific fragmentation of long-chain bases, chromatographic and metabolic alignment with synthetic references strongly supports their identity. This additional oxidative step may serve as an alternative terminal clearance route, particularly in contexts where esterification is limited.

While our study maps the overarching architecture of this novel metabolic clearance pathway, the specific identities of the downstream ω-O-acyltransferase and ω-carboxylase remain exciting frontiers for future enzymology. Potential candidates for the esterification step include enzymes with acyl-CoA:alcohol acyltransferase activity, such as DGAT1, DGAT2, or members of the MBOAT family. Similarly, identifying the specific peroxisomal or microsomal oxidases responsible for the terminal ω-carboxylation will be critical. Rather than a limitation, these unidentified enzymatic steps highlight a newly discovered, untapped regulatory landscape for atypical sphingolipids. Unlocking the specific molecular machinery behind these branches will be essential for understanding how this clearance pathway is regulated under pathological conditions.

Despite these findings, several limitations remain. Due to the complexity and novelty of the described metabolites, comprehensive sets of synthetic standards were not available. Consequently, the structural annotations for some complex downstream lipids - particularly the tritailed ceramides - rely primarily on high-resolution MS1 and MS2 fragmentation patterns and enzymatic logic, meaning their exact regiochemistry remains putative. Additionally, while we demonstrate that the initial ω-hydroxylation step attenuates acute 1-deoxySL cytotoxicity *in vitro*, the physiological bioactivity of the further downstream tritailed and carboxylated metabolites remains untested. Whether this pathway serves as an absolute detoxification route *in vivo* or if these complex lipids possess unique biological properties warrants future investigation. Finally, although we identified CYP4F2 as the primary ω-hydroxylase for 1-deoxySA, the specific enzymes responsible for the subsequent ω-O-acylation and ω-carboxylation steps have not yet been identified, leaving exciting mechanistic gaps for future enzymology studies.

In summary, our study identifies CYP4F2-mediated ω-hydroxylation as a previously undescribed mechanism for 1-deoxySL metabolic clearance and maps the metabolic conversions into distinct, less toxic downstream species. These findings expand our understanding of 1-deoxySL metabolism and provide a foundation for further investigations into their physiological and pathological significance.

## Methods

### Cell culture

HEK293 cells were grown in high-glucose Dulbecco’s Modified Eagle Medium (DMEM, Thermo Fisher Scientific) supplemented with 10 fetal bovine serum (FBS) and 1 Peniciline/Streptomycin (P/S) in a 5 CO2 incubator at 37°C. Human hepatocellular carcinoma cells (Huh-7) cells were grown in RPMI 1640 (Thermo Fisher Scientific) media were grown in high-glucose Dulbecco’s Modified Eagle Medium (DMEM, Thermo Fisher Scientific) supplemented with 10 fetal bovine serum (FBS) and 1 Peniciline/Streptomycin (P/S) in a 5 CO2 incubator at 37°C. Cells were tested for the Mycoplasma contamination.

HepaRG cells were cultured and differentiated following the method established by Gripon et. Al. (27), with minor modifications. Cells were seeded in William’s E medium supplemented with 10 fetal bovine serum (FBS), 1 GlutaMAX, 1 penicillin-streptomycin, 5 µg/mL insulin, and 50 µM hydrocortisone hemisuccinate. After an initial proliferation period of 2 weeks, differentiation was induced by supplementing the medium with 2 DMSO. The cells were maintained under these differentiation conditions for an additional 2 weeks, with media changes every 2–3 days. Differentiation status was confirmed by morphological assessment and expression of hepatic markers, as previously described. Undifferentiated control cells were maintained under identical conditions without DMSO.

### Plasmids generation

CYP4F2, CYP4F12, CYP4A11, CYP4A22 were cloned into the pcDNA3.1 Plasmid (V79020, Thermo Fisher Scientific), following manufacturer’s recommendation (Invitrogen). Generated plasmids were confirmed by Sanger sequencing.

### Overexpression of CYP isoforms

HEK293 cell lines were transfected with pcDNA3.1 Plasmid containing CYP4F2, CYP4F12, CYP4A11, CYP4A22 genes, using Lipofectamine 3000 (L3000001, Thermo Fisher Scientific). pcDNA3.1 (V79020, Thermo Fisher Scientific) was used as an Empty vector control. Transfected cells were selected by culturing in DMEM media (10 FBS) with Geneticin (G-418, Thermo Fisher Scientific, 20µg/mL) for 4 weeks.

### Stable isotope labelling assay

For the 1-deoxySL labelling assay, cells were plated at 200000 cells/mL in 6-well plates. Cells were grown for 48 hours to 70 confluency. 24 hours before harvesting, the medium was replaced with DMEM/10 FBS/1 P/S containing d_3_-1-deoxysphinganine (860474, Avanti Polar Lipids) and additional treatments (FB1, F1147, Sigma-Aldrich) as indicated in the figures. Cells were harvested by trypsinisation and counted using Beckman Coulter Z2 (Beckman Coulter). Next, cells were centrifuged at 850g at 4°C and cell pellets washed 2 times with cold PBS. Cell pellets were then frozen and kept at -20°C until further analysis.

### Supersome Enzymatic Assay

Microsomal suprasomes containing human recombinant cytochrome P450 enzymes (CYP4F2, CYP4F12, CYP4A11, and CYP3A4; Corning Gentest) were used for *in vitro* enzymatic assays. Reactions were performed in a total volume of 100 µL phosphate buffer (100 mM potassium phosphate, pH 7.4) containing 50 pmol of recombinant CYP enzyme, 1 mM NADPH, and 1 µM d_3_-1-deoxysphinganine (d_3_-1-deoxySa; Avanti Polar Lipids). Reactions were initiated by NADPH addition and incubated at 37°C for the indicated times (up to 60 minutes). Reactions were terminated by the addition of 400 µL cold methanol containing internal standards, followed by vortexing and centrifugation at 14,000 × g for 10 minutes. Supernatants were collected and subjected to high-resolution LC-MS/MS analysis to quantify ω-hydroxylated and other downstream sphingoid base metabolites.

### Cell toxicity assay

For the toxicity assay cells were grown for 72 hours in the 96-well plates in DMEM (supplemented with 10 FBS and 1 P/S) with added treatments as indicated in the figures. All conditions were corrected for the solvent concentration.

The number of viable cells was determined by quantitation of the ATP present using CellTiter-Glo **®** Luminescent Cell Viability Assay (G7570, Promega) according to the manufacturer’s recommendations. Chemiluminesce signal was collected using TECAN infinite M 200 Pro reader. Data were normalized to the average of Vehicle treated wells.

### Protein determination

For the protein content normalization, cell pellets remaining after the lipid extraction were dissolved in Urea (8M) containing 1 2-Mercaptoethanol and mixed at 800rpm at 90°C (Thermomixer (Eppendorf)) for 20 minutes. Next, protein extracts were snap frozen using dry ice, and the whole process was repeated 3 times. After centrifugation (16 100 rpm, 10 minutes), supernatants were collected and used for further analysis. Total protein amounts were determined using Bradford assay (Bio-Rad, 1:5 dilution) according to the manufacturers’ recommendation. External calibration curve of Bovine serum albumine was used to calculate total amounts of protein.

### Lipidomics analysis

Lipidomics analysis was performed as described previously (28). Briefly, extraction was performed by mixing (Thermomixer (Eppendorf), 60 minutes, 37°C) the cell pellets, plasma (100µL), or tissue homogenate with extraction buffer consisting of a mixture: methanol: methyl *tert-*butyl ether: chloroform 4:3:3 (v/v/v) and internal standards. After centrifugation (16100 rpm, 10 minutes, 37°C), the single-phase supernatant was collected, dried under N2, and stored at -20°C. Before analysis, lipids were dissolved in 100µL of MeOH Thermomixer (Eppendorf), 60 minutes, 37°C) and separated on a C30 Accucore LC column (Thermo Fisher Scientific, 150mm × 2.1 mm × 2.6 µm) or C18 ACQUITY UPLC CSH (Waters, 150mm × 2.1 mm × 1.7µm) using gradient elution with A) Acetonitrile: Water (6:4) with 10mM ammonium acetate and 0.1 formic acid and B) Isopropanol: Acetonitrile (9:1) with 10mM ammonium acetate and 0.1 formic acid at a flow rate of 260µL/minute using Transcend UHPLC pump (Thermo Fisher Scientific) at 50°C. Following gradient was used: 0min 30 (B), 0.5min 30 (B), 2min 43 (B), 3.3min 55 (B), 12min 75 (B), 18min 100 (B), 25min 100 (B), 25.5min 30 (B), 29.5min 30 (B). Eluted lipids were analysed by a Q-Exactive plus HRMS (Thermo Fisher Scientific) in positive and negative modes using heated electrospray ionisation (HESI, Shealth gas flow rate=40, Aux gas flow rate=10, Sweep gas flow rate=0, Spray voltage=3.50, Capillary temperature=320°C, S-lens RF level=50.0, Aux gas heater temperature=325°C). MS2 fragmentation spectra were recorded in data-dependent acquisition mode with top10 approach and constant collision energy 25eV. 140000 resolution was used for full MS1 and 17 500 for MS2. Peak integration was performed with TraceFinder 4.1, Skyline daily and Compound Discoverer 3.3 (Thermo Fisher Scientific). Lipids were identified by predicted mass (resolution 5ppm), retention time (RT) and specific MS2 fragmentation patterns using in house made, Lipidcreator and MSDIAL compound databases. MS2 fragmentation patterns were manually inspected using Xcalibur Qual browser. Next, lipid concentrations were normalised to the internal standards (one per class), wet weight, volume and for the cell measurements to the total protein amount.

### Long chain base analysis

Long chain bases analysis was performed as described previously (29). Briefly, 100µL of plasma was extracted using MeOH (500µL) containing internal standards. The mixture was mixed at 37°C for 60 minutes (Thermomixer (Eppendorf)) followed by centrifugation at 16 100 rpm for 10 min. Afterwards, the supernatant was collected, and 75µL of HCl (32) was added, followed by vortexing. Next, the mixture was kept at 65°C for 15 hours. Then, the mixture was neutralised using 100µL of KOH (10M) and two-phase extraction using CHCl3 and aq. NH4OH (1.5N) was performed. The CHCl3 phase was collected, dried under N2, and stored at - 20°C until further analysis. Dried-free long-chain bases were redissolved in 50µL of MeOH (70 with ten mM ammonium acetate), and 10µL was used for an LC-MS/MS analysis, using a QTRAP 6500+ (SCIEX) mass spectrometer. Peak integration was performed using Skyline daily software.

### Untargeted long chain base LC-MS/MS analysis

Free long chain bases were separated on a RP18 HPLC column (INTERCHIM UPTISPHERE C18-ODB, 125mm × 3.0mm × 3µm) using gradient elution with A.) 50 MeOH with 10mM ammonium formate and 0.2 formic acid and B.) MeOH at a flow rate of 600µL/minute using Transcend UHPLC pump (Thermo Fisher Scientific) at 50°C. The elution was carried under following gradient: 0min 35 (B), 9min 65 (B), 9.5min 100 (B), 11.50 min 100 (B), 12min 35 (B), 13min 35 (B). Eluted free long chain bases were ionised by heated electron spray ionisation (Shealth gas flow rate=55, Aux gas flow rate=15, Sweep gas flow rate=3, Spray voltage=3.50, Capillary temperature=275°C, S-lens RF level=68.0, Aux gas heater temperature=450°C) and analyzed by Q-Exactive plus HRMS (Thermo Fisher Scientific) in positive mode. MS2 fragmentation spectra were recorded in data-dependent acquisition mode with top10 approach and constant collision energy 25eV. 140000 resolution was used for full MS1 and 17500 for MS2. Peak integration was performed with Skyline daily software. Data were normalised to ISTD and total protein amount. MS2 fragmentation patterns were manually inspected using Xcalibur Qual browser.

### Study subjects for plasma collection

We obtained plasma and skin biopsy samples from a group of individuals who were recruited by Dr Pär Björklund at the Karolinska Institute in Stockholm, Sweden, during 2018-2019. This cohort was also used for research on the response-to-retention hypothesis of atherogenesis (ClinicalTrials.govID NCT03386097), which involved taking punch biopsies to measure lipid accumulation in the skin. The study was approved by the local ethics committee in Stockholm (Approval Number 2017-1942-31, 2018-1428-32, 2019-06474, 2021-06485-02) and was conducted in accordance with the Declaration of Helsinki. Plasma samples were collected at the Clinical Science Center at Karolinska University Hospital Huddinge. Participants arrived in the morning after at least 8 hours of fasting.

### Genome-wide association scan

Genotyping and imputation was described previously (30). The plasma m18:1/m18:0 ratio from 1088 participant of the CoLaus cohort has been used as a trait for a the GWAS analysis. Individuals with genotyping inconsistencies or with genotyping efficiency below 90 were removed. SNPs with genotyping efficiency below 70 or with Hardy–Weinberg p values smaller than 1E−7 were removed. Duplicate individuals and first and second degree relatives were identified by computing genomic identity-by-descent coefficients, using PLINK (42). Imputation was performed using the method of Marchini et al. (31), using IMPUTE version 0.2.0, CEU haplotypes, and the fine scale recombination map from HapMap release 21. Given the non-Gaussian distribution of the examined LCBs, inverse normal quantile transformation was applied, regressing out age, sex, and the first four ancestry principal components. The residual trait has been rescaled to have zero mean and unit variance. Linear regression analysis has been performed for 2,557,249 imputed genetic markers as explanatory variables. The probabilistic genotypes were converted to allele dosages.

### Data analysis, statistics and Figures

All the data analysis and figure preparations were performed using GraphPad Prism 9.5.1, and Excel. Statistical analysis was performed in GraphPad Prism 9.5.1. Illustrations were made using BioRender webpage (BioRender.com). ChatGPT 4o (OpenAI) was used to improve grammar and language clarity; no data were generated by the model. Final figures were prepared using Affinity Designer 2.

## Author contributions

AM performed and analyzed the experiments and drafted the manuscript. PB collected plasma samples. AJH and EY supported LC-MS/MS method development. MS performed suprasome assays and HepaRG differentiation. ES and CHA synthesized and characterized ω-OH-1-deoxySA and ω-COOH-1-deoxySA. IA, AJH, and TH critically reviewed the study and edited the manuscript.

## Supplemental data

This article contains supplemental data

