## Supplemental Data and Methods for "CYP4F2-mediated ω-hydroxylation of 1-deoxysphingolipids reveals a new hepatic detoxification pathway"

### *Supplementary Data*

#### *Synthesis of ω-carboxy-1-deoxysphinganine*

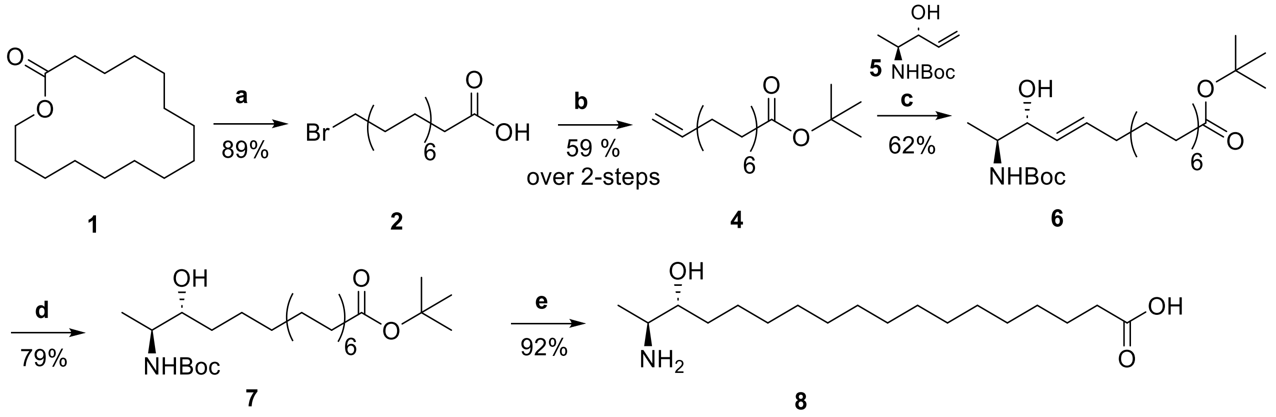


**Figure S1.** Synthesis of ω-carboxy-1-deoxysphinganine. Reagents and conditions: (i) HBr 48%, glacial acetic acid, 100˚C, 16h. (b) (i) KOtBu, THF, 78 ˚C, 16h, (ii) SOCl2, cat. DMF, 70˚C, 3h, (iii) tBuOH, NaHCO3, room temperature, 16h; (c) allyl alcohol 5 [1–3], cat. Hoveyda Grubbs II, DCM, 40˚C, 18h; (d) H2, Pt/C, MeOH, room temperature, 16h; (e) 50% TFA in DCM, 0˚C- room temperature, 4h.

The synthesis of ω-carboxy-1-deoxysphinganine was achieved via cross-coupling metathesis of allyl alcohol 5 with 14-pentadecanoic acid as a key reaction, as shown in Figure S1 (1-3). Since 14-pentadecanoic acid is not commercially available, our efforts were initially directed to synthesize 14-pentadecanoic acid starting from commercially available pentadecanolactone 1. Lactones, as cyclic esters and fatty acid precursors, undergo acid-catalyzed ring-opening hydrolysis to yield □-hydroxy acids. Thus, lactone 1 was treated with 48% HBr, followed by acetic acid to provide 15-bromopentadecanoic acid 2 in 89% yield. Subsequently, the bromo fatty acid 2 was subjected to dehydrohalogenation (elimination of HBr) using KOtBu to provide 14-pentadecenoic acid 3 in satisfactory yield. Due to the high polarity of carboxylic acids, purification by column chromatography can be challenging. To overcome this, esterification was employed to reduce polarity and improve the efficiency of chromatographic separation. Toward this, 14-pentadecenoic acid 3 was initially reacted with thionyl chloride to provide the corresponding acid chloride, which was subsequently reacted with tert-butanol in the presence of sodium hydrogen carbonate to afford tert-butyl 14-pentadecenoate 4 in 59% over the two steps. Having tert-butyl 14-pentadecenoate 4 in hand, the olefin metathesis reaction with allyl alcohol 5 was performed using Hoveyda Grubbs II catalysis to provide the protected ω-carboxy-1-deoxysphigosine 6 in a good yield (62%). The ω-carboxy-1-deoxysphigosine 6 was subsequently subjected to a hydrogenation reaction using Pt/C as a catalyst to afford the protected ω-carboxy-1-deoxysphinganine 7 in 79% yield. Finally, compound 7 was fully deprotected under acidic conditions using 50% TFA in dichloromethane to provide the desired ω-carboxy-1-deoxysphinganine 8 in

92% yield (Figure S1). The structure of the final compound was characterized by several analytical methods, including HRMS, 1H-NMR, and 13C-NMR analysis.

##### General considerations

All ragents were stored under argon unless indicated otherwise. The glassware used was dried in an oven with a magnetic stir bar inside and covered by a septum to avoid any unwanted moisture. Air-sensitive reagents were transferred via syringe under argon. The consumed solvents (e.g., DCM, ethyl ester) were distilled prior to use. Solvent removal was achieved through a rotatory evaporator equipped with a vacuum pump, controlling the pressure according to the solvent while applying 50˚C temperature for the whole removal duration to avoid any compound degradation due to extra heat. After each solvent removal, high vacuum was employed to eliminate any residual solvents. In other cases, than the ambient temperature, certain conditions were provided as: for 0˚C, water and ice combination was used; for 60˚C oil bath was used, and for -78˚C, dry ice and acetone were applied to the environment. The temperatures have been taken care of during the whole reaction time. The synthesized compounds were purified by flash column chromatography using silica gel 60 (40-63 µm). TLC analysis was achieved by TLC aluminum-backed plates, and the detection was made after soaking with staining materials such as Ninhydrin and Seebach. The structure synthesized compounds was confirmed by NMR analysis. (1H, 13C) NMR spectra of the synthesized compounds were recorded on Bruker Avance-400 and Bruker Avance-500 nuclear magnetic resonance spectrometers (1H at 400 or 500 MHz and 13C at 75 or 100 MHz) as a solution in deuterated methanol (MeOD), deuterated chloroform (CDCl3), or deuterated water (D2O) at 27°C, unless otherwise indicated.

##### Experimental synthesis and analytical data

##### Synthesis of 15-Bromopentadecanoic acid (2)

Pentadecanolactone 1 (5 g, 20.8 mmol) was dissolved in a mixture of hydrogen bromide (15 mL, HBr, 48% in water) and glacial acetic acid (125 mL) at room temperature, and the resulting reaction mixture was heated under reflux at 100 °C overnight. After completion, the mixture was allowed to cool to room temperature, and water was added to induce precipitation of the product. The resulting mixture was extracted with ethylacetate (2x 100 mL). The organic (brown) layers were separated, combined, and dried over anhydrous $Na_2SO_4$. The solvent was evaporated under reduced pressure, ensuring complete removal of residual acetic acid to afford the desired 15-bromopentadecanoic acid 2 as a pale-yellow solid (5.9 g, 18.5 mmol, 89%). The analytical data were in agreement with previously reported [4]. The compound was directly used in the next step without any further purification. 1H-NMR (400 MHz, CDCl3, ppm): δ = 3.37 (t, J= 7.2 Hz, 2H), 2.30 (t, J= 7.3 Hz, 2H), 1.85 - 1.79 (m, 2H), 1.62 - 1.58 (m, 2H), 1.44-1.38 (m, 2H), 1.25-1.20 (m, 18H). 13C-NMR (101 MHz, CDCl3, ppm): δ 179.5, 33.9, 33.6, 32.6, 29.8, 29.3, 29.0, 28.8, 28.6, 28.2, 27.9, 27.8, 27.3, 24.7.

##### Synthesis of pentadec-14-enoic acid (3)

A solution of 15-bromopentadecanoic acid 2 (3.0 g, 9.8 mmol) in dry THF (250 mL) was treated with potassium tert-butoxide (KOtBu, 1.6 g, 14.3 mmol) under an argon atmosphere. The resulting reaction mixture was allowed to stir under reflux for 16h before it was quenched at room temperature with ice-cold water. The resulting mixture was extracted with ethyl acetate (3x 100 mL), and the combined organic layers were dried over anhydrous $Na_2SO_4$. The solvent was removed under reduced pressure to afford the crude product of pentadec-14-enoic acid 3 as a yellow oil. Yield: 1.9 g (8.1 mmol, 83%). 1H-NMR (500 MHz, CDCl3, ppm) δ 5.81 (ddt, J = 17.0, 10.2, 6.7 Hz, 1H), 4.99 (ddd, J = 17.1, 3.6, 1.6 Hz, 1H), 4.92 (ddt, J = 10.2, 2.2, 1.1 Hz, 1H), 2.34 (t, J = 7.5 Hz, 4H), 2.06 – 2.01 (m, 2H), 1.66 – 1.59 (m, 4H), 1.41 – 1.25 (m, 16H). 13C-NMR (126 MHz, CDCl3, ppm) δ 180.3, 139.4, 114.2, 77.2, 61.9, 34.2, 34.0, 33.0, 29.7, 29.7, 29.7, 29.6, 29.6, 29.4, 29.3, 29.2, 29.1, 28.9, 28.3, 27.7, 24.8.

**Synthesis of tert-Butyl pentadec-14-enoate (4)**

To pentadec-14-enoic acid 3 (6.42 g, 26.7 mmol) under an argon atmosphere, thionyl chloride ($SOCl_2$, 39 mL, 53.4 mmol) was added dropwise, followed by a catalytic amount of DMF (0.5 mL). The resulting mixture was allowed to stir at 70 °C for 3 h, before it was subjected to reduced pressure under argon to remove the excess $SOCl_2$. The resulting thick mixture was dissolved in tert-butanol (tBuOH, 30 mL), and sodium bicarbonate (6.7 g, 80.1 mmol) was subsequently added under argon. The mixture was stirred overnight at room temperature and followed up by TLC analysis. After reaction completion, tBuOH was removed under reduced pressure, and the resulting mixture was suspended between water and ethylacetate. The organic layer was separated, and the aqueous layer was extracted several times with ethylacetate. The organic layers were combined, dried over anhydrous $Na_2SO_4$, and evaporated under reduced pressure. The crude product was then purified by silica gel column chromatography using a 99:1 petroleum ether:ethyl acetate (PE:EE) eluent system to provide the desired product tert-butyl pentadec-14-enoate 4, as a pale-yellow oil. Yield: 5.6 g (18.9 mmol, 71 %, 59% over 2 steps). 1H-NMR (400 MHz, CDCl3, ppm) δ = 5.81 (ddt, J = 16.9, 10.2, 6.7 Hz, 1H), 4.99 (ddd, J = 17.1, 3.6, 1.6 Hz, 1H), 4.92 (ddt, J = 10.2, 2.3, 1.2 Hz, 1H), 2.19 (t, J = 7.5 Hz, 2H), 2.03 (dt, J = 6.8, 4.4 Hz, 2H), 1.56 (dd, J = 14.1, 6.9 Hz, 2H), 1.44 (s, 9H), 1.41 – 1.34 (m, 2H), 1.33 – 1.24 (m, 16H). 13C-NMR (101 MHz, CDCl3, ppm) δ = 173.5, 139.4, 114.2, 80.0, 77.2, 35.8, 34.0, 29.7, 29.7, 29.6, 29.5, 29.3, 29.2, 29.1, 28.3, 25.3.

**Synthesis of tert-butyl (16R,17S,E)-17-((tert-butoxycarbonyl)amino)-16-hydroxyoctadec-14-enoate (6)**

To a stirred solution of allyl alcohol 5 (500 mg, 2.5 mmol) [1–3] and tert-butyl pentadec-14-enoate 4 (2.2 g, 7.5 mmol) in dry DCM under an argon atmosphere were added a catalytic amount of Hoveyda–Grubbs II catalyst was added. The resulting reaction mixture was allowed to stir at room temperature for 16h, until the allyl alcohol was consumed. The solvent was removed under reduced pressure, and the resulting residue was directly subjected to flash column chromatography using 90:10 petroleum ether:ethyl acetate (PE:EE) eluent system to provide the desired product 6 as a pale-white solid. Yield: 725 mg (1.55 mmol, 62%). 1H-NMR (400 MHz, CDCl3, ppm) δ 5.76 – 5.66 (m, 1H), 5.43 (dd, J = 15.4, 6.4 Hz, 1H), 4.72 – 4.58 (m, 1H), 4.15 – 4.07 (m, 1H), 3.77 (br.s, 1H), 2.19 (t, J = 7.5 Hz, 2H), 2.04 (q, J = 7.0 Hz, 2H), 1.61 – 1.53 (m, 2H), 1.44 (m, 16H), 1.38 – 1.21 (m, 20H), 1.07 (d, J = 6.8 Hz, 3H). 13C-NMR (101 MHz, CDCl3, ppm) δ = 173.6, 134.3, 128.5, 80.1, 77.2, 35.9, 32.6, 29.8, 29.7, 29.5, 29.4, 29.3, 28.6, 28.4, 25.4, 15.8.

**Synthesis of tert-butyl (16R,17S,E)-17-((tert-butoxycarbonyl)amino)-16-hydroxyoctadec-14-enoate (7)**

A stirred solution of compound 6 (550 mg, 1.2 mmol) in a dry mixture of THF and methanol (1:1, 15 mL) under an argon atmosphere at room temperature was subjected to a catalytic amount of 10% Pt/C. The argon was removed using a vacuum pump, and the mixture was connected to a hydrogen balloon. The resulting mixture was stirred under the same conditions for 16h until HPLC-MS analysis revealed complete reduction. The reaction mixture was subsequently filtered through a pad of Celite, and the filtrate was concentrated under reduced pressure. The obtained residue was purified by flash column chromatography using an 85:15 petroleum ether:ethyl acetate (PE:EE) eluent system to provide compound 7 as a waxy solid. Yield: 445 mg (0.95 mmol, 79%). 1H-NMR (400 MHz, CDCl3, ppm) δ = 3.72 – 3.60 (m, 2H), 2.19 (t, J = 7.5 Hz, 2H), 1.62 – 1.53 (m, 2H), 1.44 (m, 18H), 1.42 – 1.35 (m, 2H), 1.32 – 1.22 (m, 22H), 1.07 (d, J = 6.8 Hz, 3H) ppm. 13C-NMR (101 MHz, CDCl3, ppm) δ 173.5, 156.0, 77.2, 35.8, 33.6, 29.8, 29.7, 29.6, 29.4, 29.2, 28.5, 28.3, 26.2, 25.3, 14.5.

**Synthesis of (16R,17S)-17-amino-16-hydroxyoctadecanoic acid (8)**

A stirred solution of compound 7 (300 mg, 0.64 mmol) in dry DCM (5 mL) at 0°C was treated with trifluoracetic acid (5 mL). The resulting reaction mixture was allowed to stir under the same conditions for 1h, and an additional 3h at room temperature, before the solvent was removed under reduced pressure. The resulting residue was dissolved in DCM and washed with sat. sodium hydrogen carbonate solution, brine, dried over anhydrous $Na_2SO_4$, and evaporated under reduced pressure to provide the desired product as a white solid. Yield: 185 mg (0.59 mmol, 92%). 1H-NMR (400 MHz, MeOD, ppm) δ = 3.72 – 3.66 (m, 1H), 3.26 (dd, J = 6.7, 2.8 Hz, 1H), 2.28 (t, J = 7.4 Hz, 2H), 1.64 – 1.55 (m, 2H), 1.49 – 1.40 (m, 2H), 1.39 – 1.27 (m, 22H), 1.21 (d, J = 6.8 Hz, 3H). 13C-NMR (101 MHz, MeOD, ppm) δ = 71.7, 52.6, 49.6, 49.4, 49.2, 49.1, 49.0, 48.8, 48.6, 48.4, 34.9, 34.0, 30.8, 30.7, 30.6, 30.4, 30.2, 27.0, 26.1, 12.0.

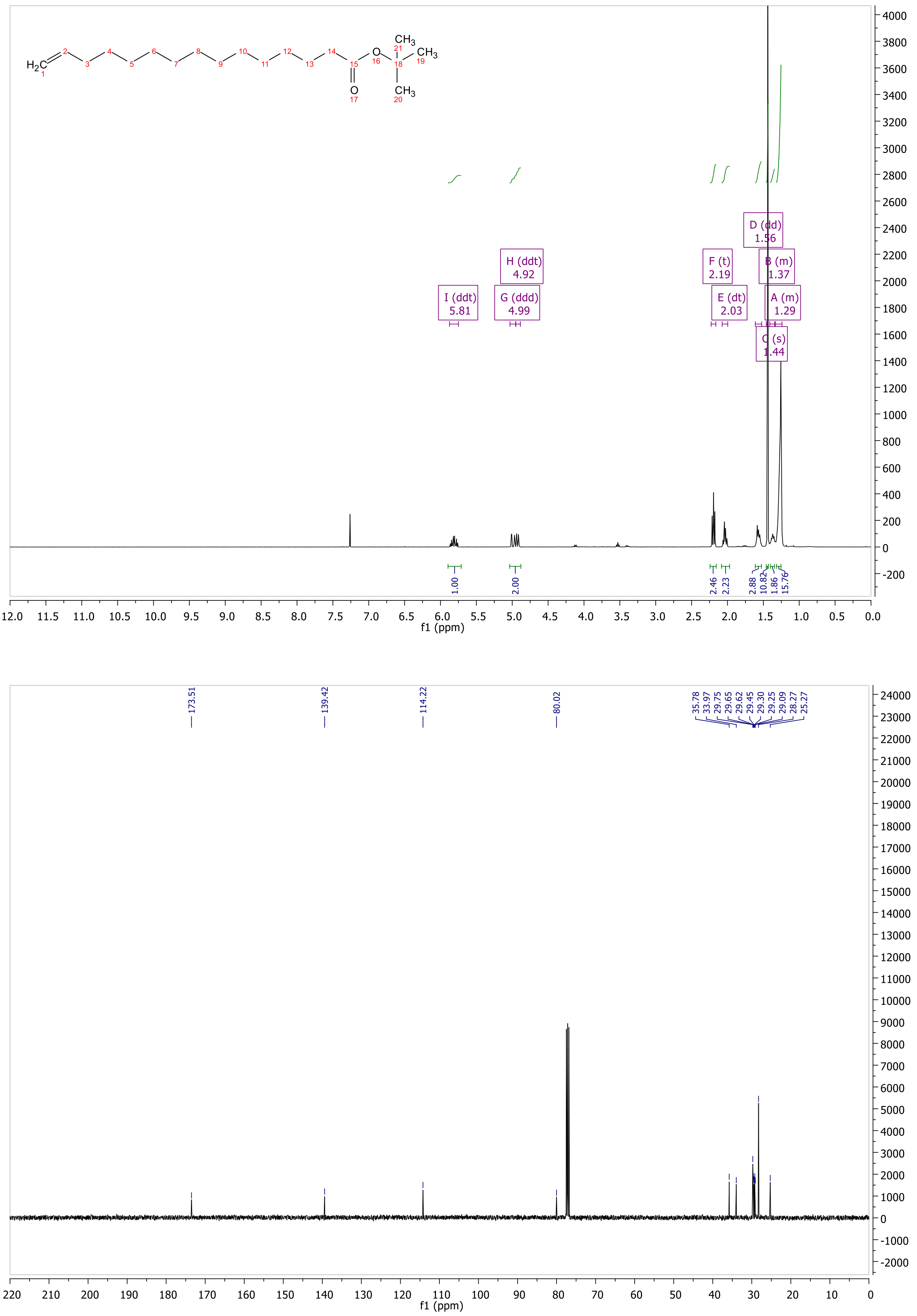


**Figure S2.** $^{1}H$-NMR and $^{13}C$-NMR of compound **4**.

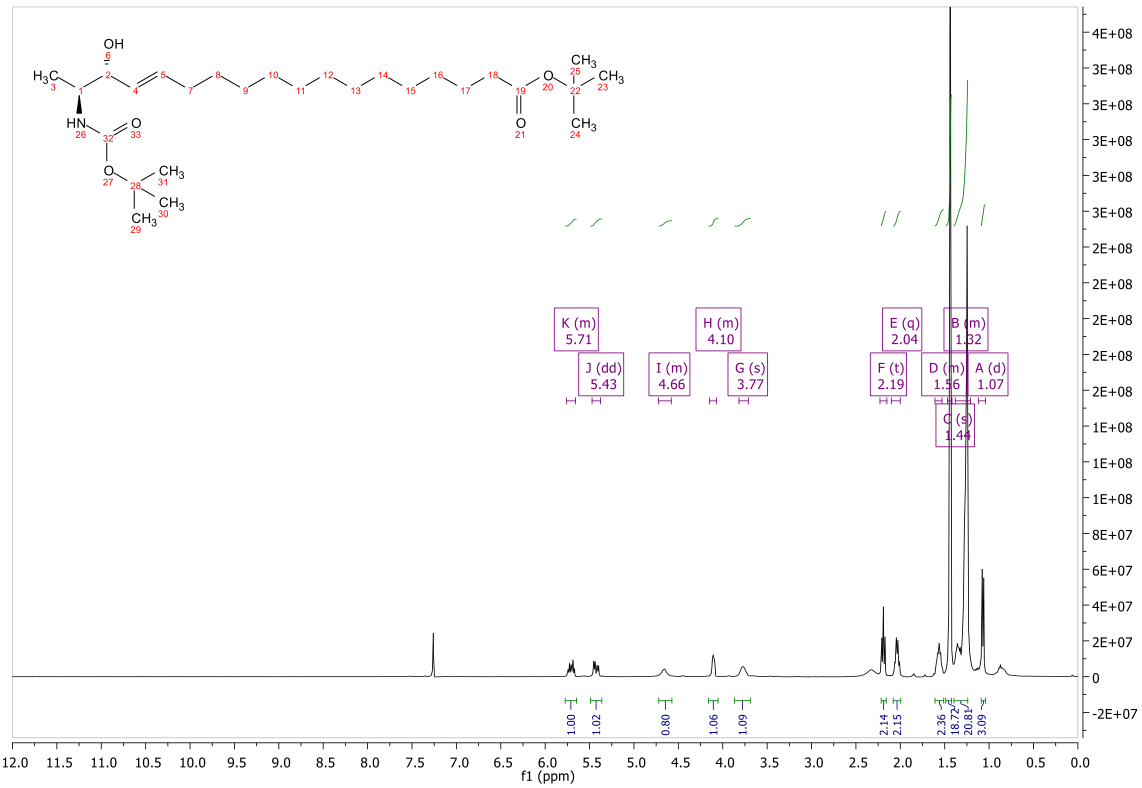


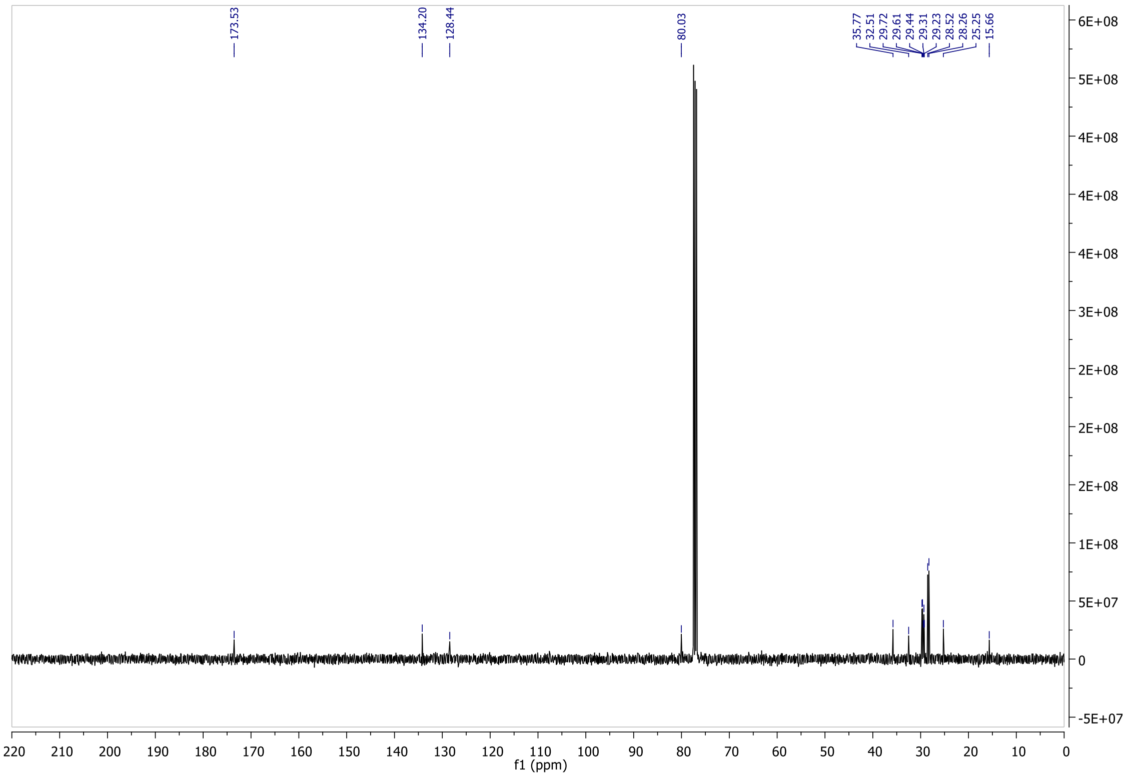


**Figure S3.** $^{1}H$-NMR and $^{13}C$-NMR of compound **6**.

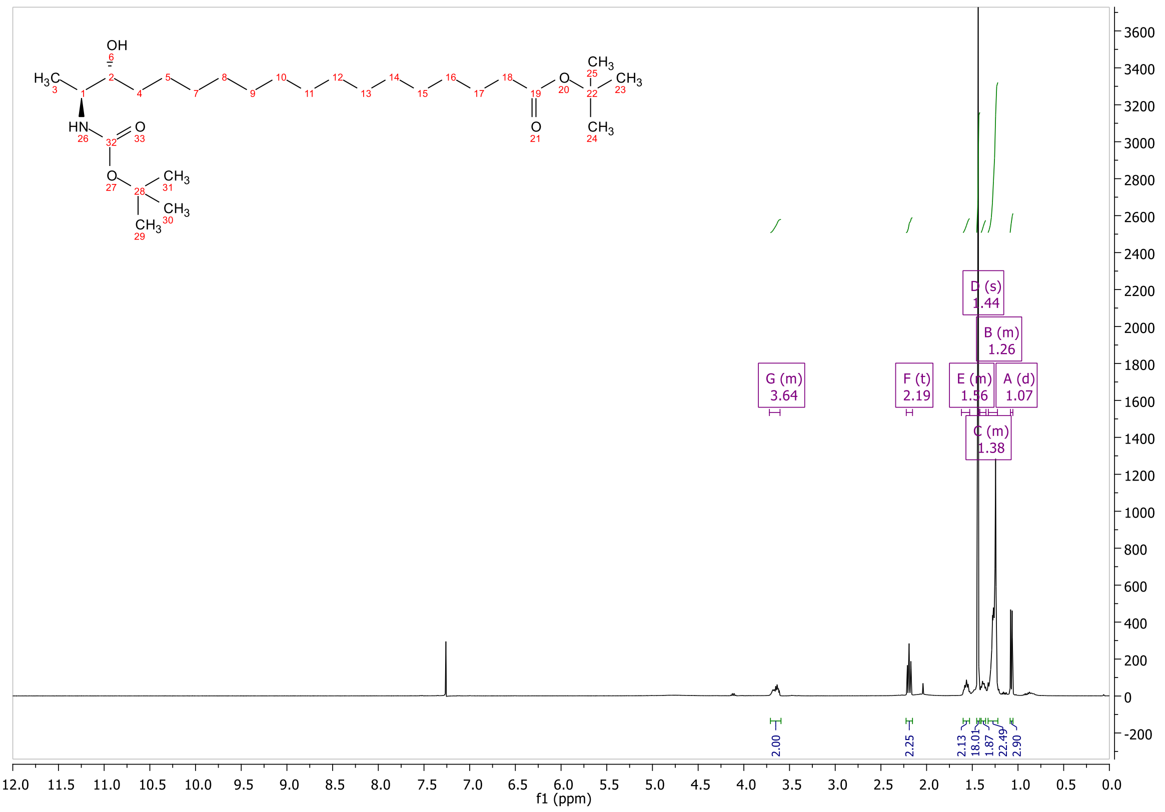

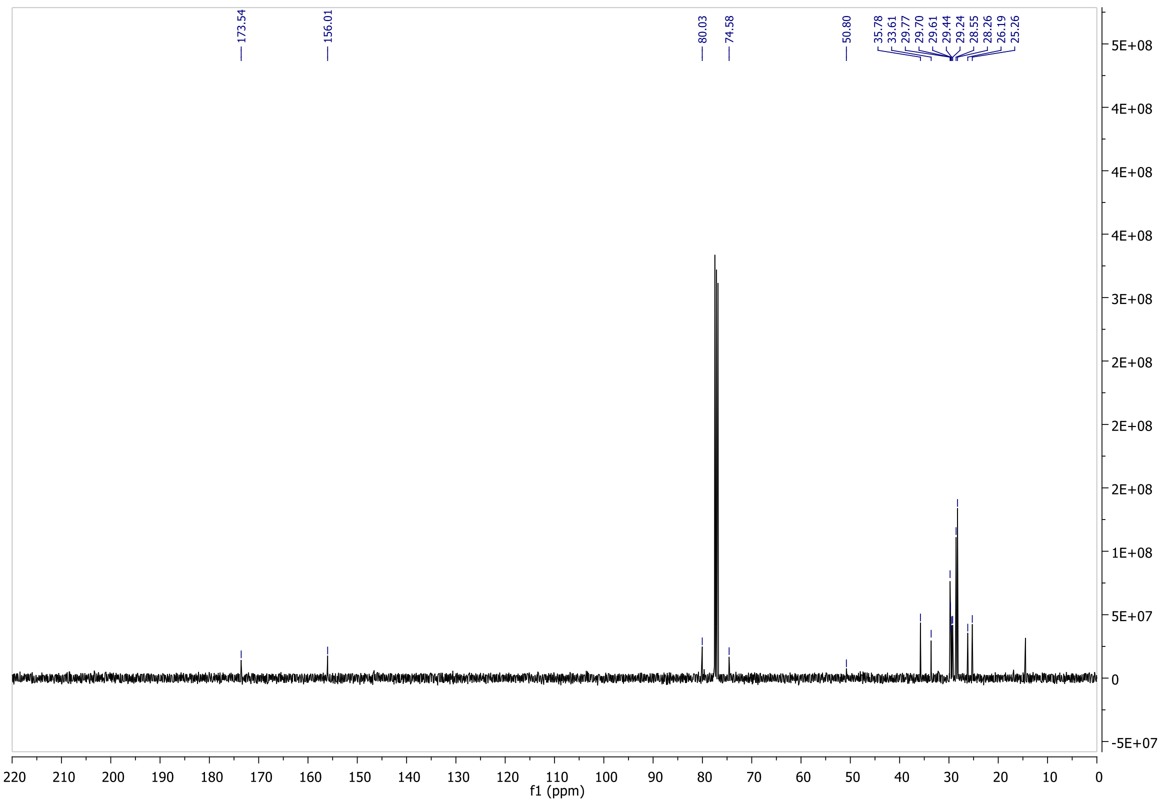

**Figure S4.** $^{1}$H-NMR and $^{13}$C-NMR of compound **7**.

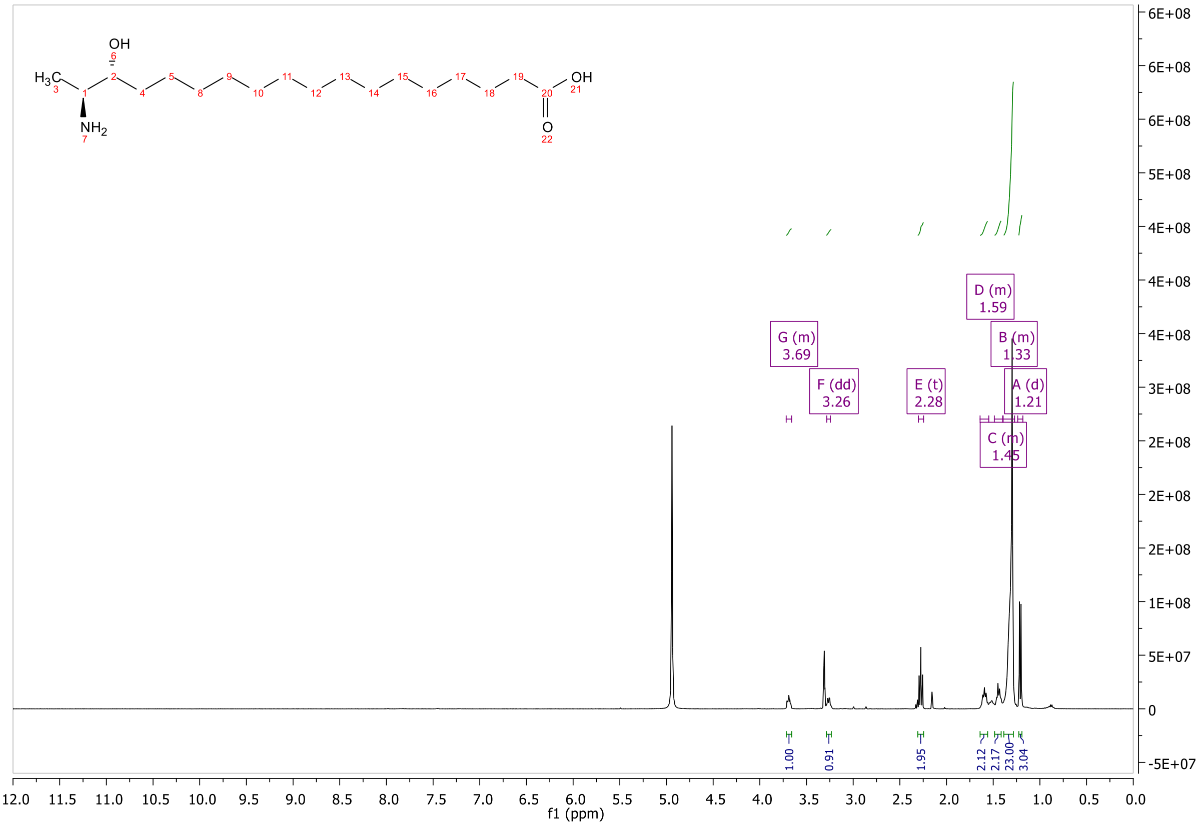


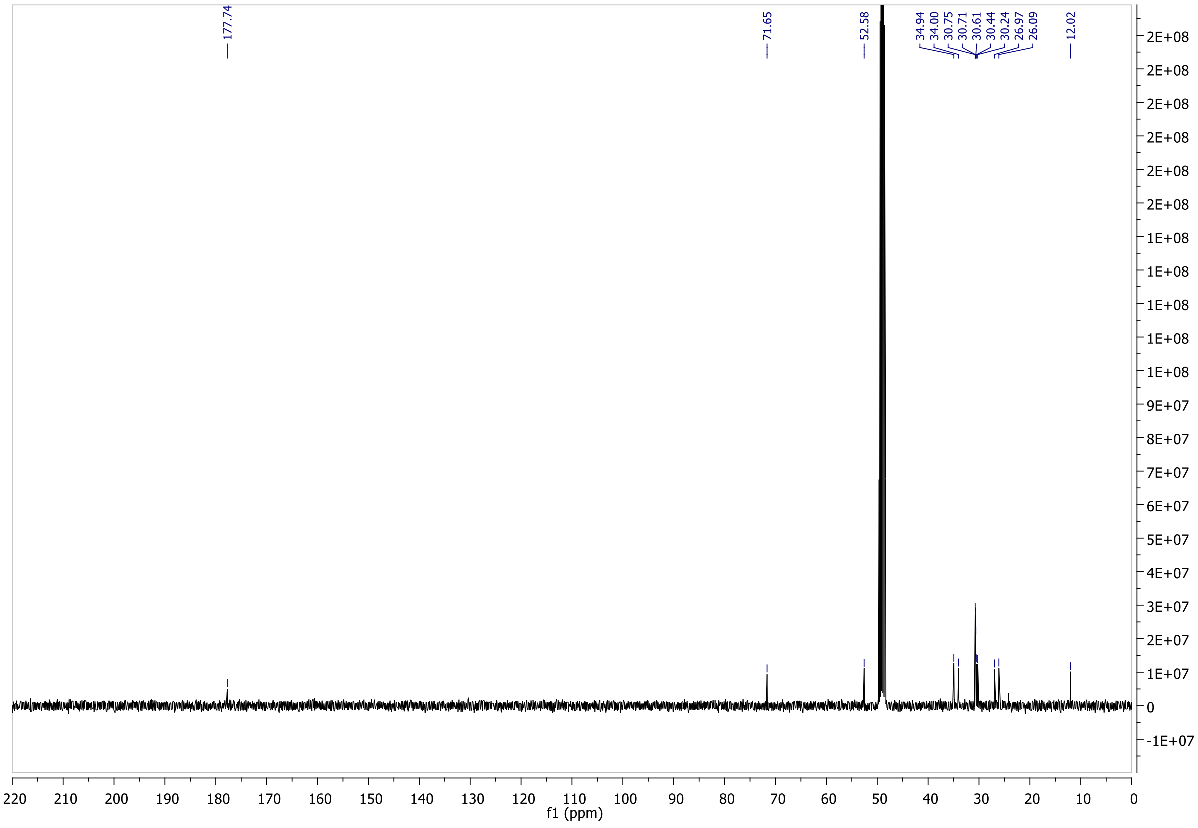


**Figure S5.** ¹H-NMR and ¹³C-NMR of compound **8**.

#### *Supplementary References*

1. Saied, E. M., T. L. S. Le, T. Hornemann, and C. Arenz. 2018. Synthesis and characterization of some atypical sphingoid bases. *Bioorgan Med Chem* **26**: 4047–4057.
2. Saied, E. M., S. Banhart, S. E. Burkle, D. Heuer, and C. Arenz. 2015. A series of ceramide analogs modified at the 1-position with potent activity against the intracellular growth of Chlamydia trachomatis. *Future Med Chem* **7**: 1971–1980.
3. Saied, E. M., and C. Arenz. 2021. Stereoselective Synthesis of Novel Sphingoid Bases Utilized for Exploring the Secrets of Sphinx. *Int J Mol Sci* **22**.

***List of internal standards used for lipidomics analysis:***

D5-1-desoxymethylsphinganine (m17:0, 860476, Avanti Polar Lipids) 100pmol/sample
D7-Sphinganine (d18:0, 860658, Avanti Polar Lipids) 100pmol/sample
D7-Sphingosine (d18:1, 860657, Avanti Polar Lipids) 100pmol/sample
D7-Sphingosine-1-phosphate (d18:1, 860659, Avanti Polar Lipids) 50pmol/sample
1-deoxydihydroceramide (m18:0/12:0, 860460P, Avanti Polar Lipids) 100pmol/sample
1-deoxyceramide (m18:1/12:0, 860455, Avanti Polar Lipids) 100pmol/sample
Dihydroceramide (d18:0/12:0, 860635, Avanti Polar Lipids) 100pmol/sample
Ceramide (d18:1/12:0, 860512, Avanti Polar Lipids) 100pmol/sample
SM (d18:1/12:0, 860583, Avanti Polar Lipids) 100pmol/sample
Glucosylceramide (d18:1/8:0, 860540, Avanti Polar Lipids) 100pmol/sample
SPLASH lipidomics standard (330707, Avanti Polar Lipids) 2.5μL/sample

Transitions used for the identification of Sphingolipids and 1-deoxySphingolipids:
1-deoxySL:
$[M+H]^+ \rightarrow [M+H - H_2O]^+$ , $[M+H]^+ \rightarrow [M+H - H_2O - FA]^+$
Ceramides and dihydroCeramides:
$[M+H]^+ \rightarrow [M+H - H_2O]^+$ , $[M+H]^+ \rightarrow [M+H - H_2O - FA]^+$ ,
$[M+H]^+ \rightarrow [M+H - 2xH_2O - FA]^+$
HexosylCeramides:
$[M+H]^+ \rightarrow [M+H - Hexosyl]^+$ , $[M+H]^+ \rightarrow [M+H - H_2O - FA - Hexosyl]^+$ ,
$[M+H]^+ \rightarrow [M+H - 2xH_2O - FA - Hexosyl]^+$
Sphingomyelins
$[M+H]^+ \rightarrow [PO4\text{-}Choline]^+$ , $[M+H]^+ \rightarrow [M+H - H_2O - FA - PO4\text{-}Choline]^+$ ,
$[M+H]^+ \rightarrow [M+H - 2xH_2O - FA - PO4\text{-}Choline]^+$
Sphingosine-1-phosphate
$[M+H]^+ \rightarrow [M+H - 2xH_2O - PO4]^+$, $[M+H]^+ \rightarrow [M+H - H_2O - PO4]^+$
Free canonical long chain bases (Sphingosine and Sphinganine):
$[M+H]^+ \rightarrow [M+H - H_2O]^+$, $[M+H]^+ \rightarrow [M+H - 2xH_2O]^+$
Free atypical long chain bases
$[M+H]^+ \rightarrow [M+H - H_2O]^+$

Note: FA represents corresponding fatty acyl.

***List of internal standards used for long chain base analysis:***

D5-1-desoxymethylsphinganine (m17:0, 860476, Avanti Polar Lipids) 200pmol/sample
D7-Sphinganine (d18:0, 860658, Avanti Polar Lipids) 200pmol/sample
D7-Sphingosine (d18:1, 860657, Avanti Polar Lipids) 200pmol/sample
